# Gradations in protein dynamics captured by experimental NMR are not well represented by AlphaFold2 models and other computational metrics

**DOI:** 10.1101/2024.07.17.603933

**Authors:** Jose Gavalda-Garcia, Bhawna Dixit, Adrián Díaz, An Ghysels, Wim Vranken

**Author notes:** Authors contributed equally to this work (Jose Gavalda-Garcia, Bhawna Dixit).

## Abstract

The advent of accurate methods to predict the fold of proteins initiated by AlphaFold2 is rapidly changing our understanding of proteins and helping their design. However, these methods are mainly trained on protein structures determined with X-ray diffraction, where the protein is packed in crystals at often cryogenic temperatures. They can therefore only reliably cover well-folded parts of proteins that experience few, if any, conformational changes. Experimentally, solution nuclear magnetic resonance (NMR) is the experimental method of choice to gain insight into protein dynamics at near physiological conditions. Computationally, methods such as molecular dynamics and Normal Mode Analysis (NMA) allow the estimation of a protein’s intrinsic flexibility based on a single protein structure. This work addresses, on a large scale, the relationships for proteins between the AlphaFold2 pLDDT metric, the observed dynamics in solution from NMR metrics, interpreted MD simulations, and the computed dynamics with NMA from single AlphaFold2 models and NMR ensembles. We observe that these metrics agree well for rigid residues that adopt a single well-defined conformation, which are clearly distinct from residues that exhibit dynamic behavior and adopt multiple conformations. This direct order/disorder categorisation is reflected in the correlations observed between the parameters, but becomes very limited when considering only the likely dynamic residues. The gradations of dynamics observed by NMR in flexible protein regions are therefore not represented by these computational approaches. Our results are interactively available for each protein from https://bio2byte.be/af_nmr_nma/.

## 1 Introduction

The advent of accurate methods to predict the fold of proteins initiated by AlphaFold2 [1] is rapidly changing our understanding of proteins [2, 3] and helping their design [4–6]. However, these methods only reliably cover well-folded parts of proteins that experience few, if any, conformational changes. Methods are now being developed that predict multiple conformations, for example by modifying the multiple sequence alignment input that captures evolutionary information [7] or by adapting the underlying deep learning model [8]. Unfortunately, reliable data on such multiple conformations that enables validation of these methods remains scarce. For example, it is estimated that up to 4% percent of proteins can change their fold, while we only have experimentally detected very few of such changes [9]. In addition, smaller conformational changes, which can be functionally important, are difficult to detect and often only captured by specific experimental NMR relaxation measurements[10]. Finally, protein dynamics, which encompass the timescales of interconversion between multiple conformations, are functionally highly relevant but rarely well-defined experimentally, with computational developments essential [11]. While computational successes are reported on individual protein cases, an extensive large scale comparison for proteins between predicted models and their computed dynamics to experimental observations of dynamics and the gradations thereof is still missing.

AlphaFold2 captures the local accuracy of the structure predictions by the predicted local distance difference test (pLDDT) [1, 12]. The pLDDT value for a residue is an estimation of the resemblance between the prediction and an experimentally determined protein structure [13]. Given that the dataset employed to train AlphaFold2 exclusively contains static protein structures [1] obtained with experimental X-ray diffraction and cryo-electron microscopy (cryo-EM), high pLDDT values therefore indicate that a protein residue is likely in a well-folded rigid state that can be represented by fixed coordinates [14]. Meanwhile, longer loops are often missing in the X-ray diffraction structures. This is illustrated by AlphaFold2’s capacity to predict “disorder” versus “order” for protein residues from pLDDT values [15]. However, such a “disorder” versus “order” distinction does not capture gradations in dynamics, only its presence or absence. Indeed, cryo-EM and, typically, X-ray diffraction study proteins at cryogenic temperatures, with proteins in crystalline form in diffraction studies. These measurements do thus not represent the native dynamics and multiple conformations that proteins experience in solution at temperatures enabling life [16]. Such dynamic residues cannot be described by a static set of coordinates and often will be poorly resolved, or might even prevent the formation of crystals altogether [17]. Instead, proteins with multiple conformations, for example as observed for intrinsically disordered regions (IDRs) and proteins (IDPs), require ensemble representations of structures [18] and other descriptors for their dynamics at different timescales.

The experimental method of choice to gain insight into protein dynamics is solution nuclear magnetic resonance (NMR), which can study protein dynamics and allows structural space at near physiological conditions [17]. Dedicated NMR experiments can accurately determine protein dynamics at different timescales, often captured by the *S*^2^ order parameter [19], but such measurements remain scarce. More readily available are NMR chemical shift values [20] stored at the BioMagResBank (BMRB). Chemical shifts can be interpreted to gain less accurate, but still useful, information on dynamics and conformation at the per-residue level. Notably, the random coil index (RCI) estimates the backbone dynamics at the residue level from a simple model that interprets experimental chemical shift values. It also provides an approximation of the backbone *S*^2^ order parameter as the 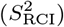 value [21, 22]. Additionally, ShiftCrypt [23] presents a machine-learning based alternative by deriving a single encoded per-residue value from NMR chemical shifts, which captures a combination of conformational preference and dynamics at the protein residue level. These residue-level experimental metrics therefore allow a direct comparison of experimentally determined protein dynamics, of movements averaged when faster than ms timescales, with structure-related per-residue values such as pLDDT. The use of RCI in this context was previously illustrated by an investigation of predicted AlphaFold2 models versus NMR models calculated from experimental data. The ANSURR method (Accuracy of NMR Structures Using random coil index (RCI) and Rigidity) used a dataset of 904 human proteins [24] to compare their scaled RCI value to a local structure-based rigidity measure [25]. The NMR models with highest-scoring ANSURR scores in each ensemble, indicating a good match between in-solution observations and structure models, showed accuracy comparable with AlphaFold2, with AlphaFold2 performing significantly better in 30% of cases, particularly in relation to regions with extensive hydrogen-bond (H-bond) networks. Only in 2% of cases, the ANSURR score of the NMR structure ensemble was higher, primarily in dynamic regions [24], indicating that AlphaFold2 struggles with these regions.

Computationally, simulations of a 3D representation of the protein can also provide valuable insights on their dynamics. With molecular dynamics (MD) simulations, the protein’s flexibility and conformational states can be investigated. Besides MD, Normal Mode Analysis (NMA) allows the estimation of a protein’s intrinsic flexibility based on a single protein structure [26]. NMA provides information on the low-energetic motions that are accessible to the system at finite temperature. These large-amplitude motions are often related to biological function. Usually the elastic network model (ENM) approximation is applied, which models the protein as a set of C_α_ beads in 3D space interacting with each other within a certain cutoff range [27]. The ENM therefore successfully captures the connectivity of the protein, which is a primary factor for the protein’s flexibility. Moreover, NMA is computationally orders of magnitude cheaper than a standard MD simulation, and hence it is a valuable tool to swiftly assess possible dynamics for an extended dataset of proteins, especially since the low-frequency normal modes generated by NMA have been shown to effectively capture the collective motions of proteins observed in both NMR experiments and MD simulations [28–31]. This correlation between NMA predictions and experimental observations highlights the significant influence of the backbone in describing collective dynamics, and NMA can therefore in principle be used to assess the flexibility of static AlphaFold2 models.

This work addresses, on a large scale, the relationships for proteins between the AlphaFold2 pLDDT metric, the observed dynamics in solution from NMR metrics, interpreted MD simulations and the computed dynamics with NMA from single AlphaFold2 models and NMR ensembles. We observe that these metrics agree well for rigid residues that adopt a single well-defined conformation, which are clearly distinct from residues that exhibit dynamic behavior and adopt multiple conformations. This direct order/disorder categorisation is reflected in the correlations observed between the parameters, but becomes very limited when considering only the likely dynamic residues. The gradations of dynamics observed by NMR in flexible protein regions are therefore not represented by these computational approaches. Our results are interactively available for each protein from https://bio2byte.be/af_nmr_nma/.

## 2 Methods

### 2.1 Overview of information in three datasets

Three datasets were constructed: the 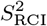 dataset, the *S*^2^ dataset, and the Molecular Dynamics (MD) dataset (Table 1). Besides common elements, such as the the protein sequence information, the datasets differ by per-residue metrics such as 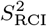 values. Per dataset, all the per-residue information for a given entry is collected in integrated Pandas data frames [32, 33] that are made available on https://zenodo.org/doi/10.5281/zenodo.10977724.

**Table 1:**
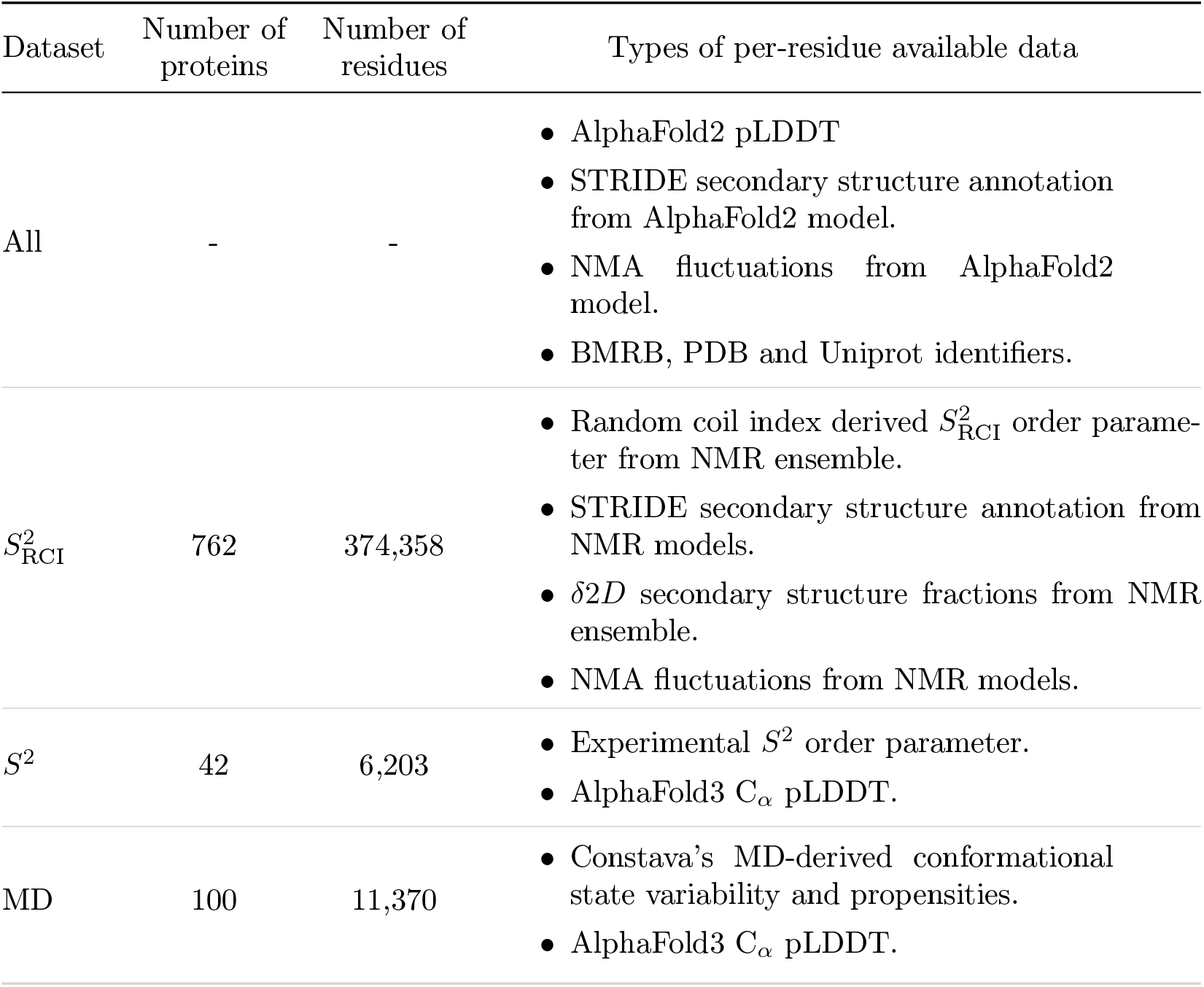
Dataset overview. Content of the 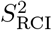, *S*^2^ and Molecular Dynamics (MD) datasets used in this work. The full list of data elements can be found in the supplementary data frames (https://zenodo.org/doi/10.5281/zenodo.10977724).

#### 2.1.1 Structure models and NMR data

##### AlphaFold2 structures

were downloaded from AlphaFold2’s EBI database [13] (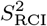 dataset) or generated on the Vlaams Supercomputer Centrum (VSC) infrastructure (*S*^2^ and MD datasets). Downloads from the EBI database [13] were performed with the in-house software package AlphaFetcher (https://pypi.org/project/AlphaFetcher/). Other AlphaFold2 structures were calculated on the VSC, with the cut-off date for the structures employed as templates was the 15^th^ of February 2021, for uniformity with the 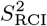 dataset. In-house calculations of AlphaFold2 structures ensured that the dataset was not reduced by the overlap between our dataset and AlphaFold2’s database. AlphaFold3 C_α_ pLDDT values were calculated on the publicly available AlphaFold3 server https://golgi.sandbox.google.com/, excluding the 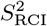 dataset entries due to current job number limitations on this server, restricting scalability. The AlphaFold2 and/or AlphaFold3 pLDDT values are per-residue values and are integrated in the Pandas data frame [32, 33].

##### Chemical shift data

was collected from the Biological Magnetic Resonance Data Bank (BMRB) [34], using previously described criteria [35, 36]. Briefly, only entries between reported pH 5–7, temperature 293–313K, for which chemical shift data was available for ^1^H, ^13^C and ^15^N for at least half of the residues in the sequence were selected. Entries with samples containing agents that strongly influence protein behaviour (Supplementary Table 1 in [35]) were excluded, and chemical shift re-referencing was performed with VASCO [37].

##### NMR structure ensembles

corresponding to the BMRB sequences were collected from the Protein Data Bank (PDB) [38] only if there was a 100% sequence identity match between the BMRB and PDB sequence. For this set of proteins, the AlphaFold2 database was queried [13]. If the AlphaFold2 structure was available, the protein was withheld, and the BMRB and AlphaFold2 sequences were locally aligned. Only entries where the longest uninterrupted aligned fragment covered at least 40% of the BMRB sequence were finally retained, resulting in a final dataset of 762 proteins. The 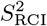 dataset has thus 762 protein entries with each an AlphaFold2 model and each an NMR ensemble. As many proteins have more than 1 NMR model, the total number of NMR models is 14,334.

##### Experimental *S*^2^ values

were collected from the BMRB for a set of 52 proteins. Of those, 42 proteins were retained to build the *S*^2^ dataset, after removing overlap from multiple BMRB IDs mapping to the same Uniprot accession code (Table. 1): in these cases, the BMRB entry with the highest number of *S*^2^ data points was retained. The experimental *S*^2^ order parameters of a protein are per-residue values and are integrated in the Pandas data frame for each of the 42 entries [32, 33].

#### 2.1.2 Information derived from models and NMR data

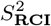 **order parameters**

were calculated for these 762 proteins from the processed chemical shift values using the RCI software [22] and integrated in the Pandas data frame for each entry as per-residue values [32, 33].

##### STRIDE secondary structure

assignments for all residues using STRIDE [39, 40] were obtained for all available AlphaFold2 models and for all models in each available NMR ensemble. Moreover, for proteins with multiple NMR structures in their ensemble, the “STRIDE unique” value of a residue indicates that the STRIDE secondary structure of that residue is the same for all models in the NMR ensemble, whereas the “STRIDE consensus” indicates the majority secondary structure assignment along all models. The STRIDE secondary structure assignments are per-residue values and are integrated in the Pandas data frame [32, 33].

##### MD simulations

of single chain entries from the Constava dataset [36] were employed for this work, totalling 100 protein ensembles. The **conformational state variability (Constava)** was determined according to the described methodology [36]. Trajectories were sampled using window size 3 sampling, a recommended approach for balanced variability computation for time-series data (such as the MD simulations of this dataset). All calculations were performed utilizing the PyPI package associated with the Constava method with its quick grid interpolation model https://pypi.org/project/constava/. The Constava values of a protein are per-residue values and are integrated in the Pandas data frame for each entry [32, 33].

#### 2.1.3 Normal Mode Analysis

Root mean square fluctuations (RMSF) from NMA were calculated on the AlfphaFold2 structures for all datasets and on the NMR ensembles for the 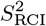 dataset.

The normal modes and eigenfrequencies of all AlphaFold2 and NMR ensemble models were computed with the open-source WEBnma webserver [27, 41]. Based on the normal modes with lowest eigenvalues, the atoms’ fluctuations were estimated under thermal equilibrium [42]. The squared fluctuation of atom *i* is

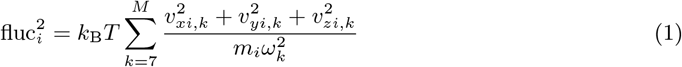

Here, *k*_B_ is the Boltzmann constant, *T* is temperature (here 300 K), *M* − 6 the number of contributing eigenvectors, *m*_*i*_ is the mass of the *i*^th^ amino acid residue, 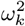 is the *k*^th^ eigenvalue, *v*_*xi,k*_ is the Cartesian *x*-component for C_α_ atom *i* in the corresponding *k*^th^ normal mode vector, and similarly for *v*_*yi,k*_ and *v*_*zi,k*_. The normal mode vectors *v*_*k*_ are mass-weighted and normalized [42]. The sum skips the 6 zero-frequency modes corresponding to global translation and rotations. In practice, only the lowest 200 non-trivial normal mode vectors (*M* = 206) were included in the sum. Based on these C_α_ atom fluctuations, the root mean square fluctuations (RMSF)_i_ of each residue *i*, which is a measure of the average fluctuation or displacement of individual atoms from their mean positions, was obtained by taking the square root,

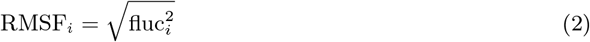

The normalized eigenvectors and eigenvalues from WEBnma were used to construct the RMSF profile with Eqs. 1-2.

The standard WEBnma calculations led to several unrealistically large RMSF values in unfolded parts of the protein, which are too loosely connected by the elastic network model. These regions are often present at the termini of the protein. Therefore, a pre-processing step was carried out, as detailed in the Supplementary Information (Section Truncation criterion). In summary, the preprocessing step involved truncating the Nand/or the C-terminal residues. To do so, the number of C_α_ contacts within 10 Å of each C_α_ were computed using MDAnalysis. If terminal residues with fewer than 13 contacts were present, these were removed until the first residue with at least 13 contacts (Supplementary Fig. 14). Unfolded terminal residues are thus removed, while highly-connected terminal residues are not (Supplementary Fig. 15). The resulting truncated proteins are only used for the calculation of the RMSF based on the C_α_ atoms. To do so, WEBnma was re-run on these truncated structures and the per-residue RMSF computed with Eqs. 1-2.

For the 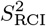 dataset, the NMA fluctuations were computed with WEBnma for the 762 AlphaFold2 models using the truncation approach. The truncation criteria led to 755 truncated proteins, while 7 did not require cutting of termini. The NMA fluctuations were also computed with WEBnma on each model of the 762 NMR ensembles of the 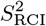 dataset. Out of the 14,334 3D models in the NMR ensembles, some models had to be discarded. One protein did not show an overlap after truncation between the AlphaFold model and its NMR models (10 discarded); two protein pdb-files could not be split (2 discarded); 13 proteins produced a WEBnma error due to invalid distance (<0.278 Å) between C_α_ atoms for cis peptide bonds (174 discarded). Moreover, this WEBnma error occurred in some isolated NMR models of the other proteins (51 discarded), however without removing entirely the protein entry as it had at least one successful WEBnma calculation. Thus, in this paper, RMSF will be reported for 762 proteins using the NMA fluctuations of 762 AlphaFold2 models, as well as for a subset of 746 proteins using the NMA fluctuations of 14,069 NMR models.

For the *S*^2^ dataset and MD dataset, the RMSF was successfully computed with WEBnma using the truncation approach for all 42 (41 needed truncation) and 100 (30 needed truncation) AlphaFold2 structures, respectively.

The RMSF are per-residue values and are integrated in the Pandas data frame for each protein entry [32, 33]. As a protein can have multiple NMR models, the entry may contain multiple RMSF profiles corresponding to different models in its NMR ensemble.

#### 2.1.4 Normalisation of pLDDT ranges

To elucidate the propensity of each pLDDT range towards each conformational state, we utilized the following normalization process:

- Let *S* represent a specific secondary structure of a residue.
- The pLDDT ranges are categorized into low, mid, and high, corresponding to the low, mid, and high values, respectively.
- *N*_*S*,low_, *N*_*S*,mid_, and *N*_*S*,high_ denote the counts of residues in the secondary structure *S* that reside within the specified pLDDT ranges.
- The total count of residues pertaining to structure *S* across all pLDDT ranges is expressed as

*T*_S_ = *N*_S,low_ + *N*_S,mid_ + *N*_S,high_.

The normalized counts for residues in secondary structure *S* across different pLDDT ranges (low, mid and high) are calculated using the formula:

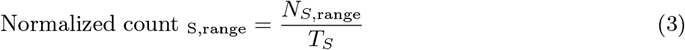

Following this, the relative normalized fractions are calculated using the formula:

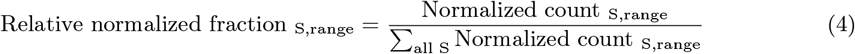

which ensures the sum over fractions equals 1 within each pLDDT range. These two steps convert the raw counts into proportional measures that reflect the relative abundance of each secondary structure *S* within each pLDDT range.

### 2.2 Data analysis and plotting

A docker file containing all necessary code for the analysis and plots in this work can be found in https://hub.docker.com/repository/docker/jgavalda/alphafold_analysis_pipeline/. The code (available on https://bitbucket.org/bio2byte/af2_analysis_datagen_and_plots/src/main/) contains all necessary tools to download AlphaFold2 structures and integrate them with the different data sources. The generated AlphaFold2 and AlphaFold3, as well as the NMA calculations are not included in this pipeline to alleviate the required computing resources. The complete pipeline can be ran in approximately 2 hours with a standard desktop computer.

## 3 Results

This study investigates to which extent AlphaFold2 models and the associated pLDDT metric can capture protein dynamics information via three datasets: the NMR chemical shift derived 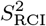, the NMR directly measured *S*^2^ order parameter and molecular dynamics (MD) simulations leading to traditional MD metrics and the conformational state variability (described in Methods 2.1 and Table 1).

### 3.1 Secondary structure comparison between AlphaFold2 models and NMR ensembles

NMR structure ensembles are calculated from available experimental NMR information such as NOEs from NOESY spectra, which are converted to distances used in the NMR structure calculations. NMR structure ensembles therefore capture experimental NMR information, but only to a certain degree as the data to model process is highly complex [43]. A comparison between predicted AlphaFold2 models and experimentally derived NMR structure ensembles is therefore a relevant first step to detect discrepancies. We focused on the STRIDE [39, 40] secondary structure assignment for AlphaFold2 models and the corresponding NMR ensembles in the 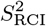 dataset (see Methods 2.1.1). The STRIDE assignments from the AlphaFold2 structures were stratified according to their pLDDT ranges in high (> 80), mid (80 > pLDDT > 60) and low (< 60). The abundance of residues belonging to each secondary structure in each of these ranges was normalised (see Methods 2.1.2) to reflect the tendencies of each secondary structure within each pLDDT range. This enables a comparison between the AlphaFold2 and NMR secondary structure (Fig. 1 & Supplementary Fig. 2).

**Figure 1:**
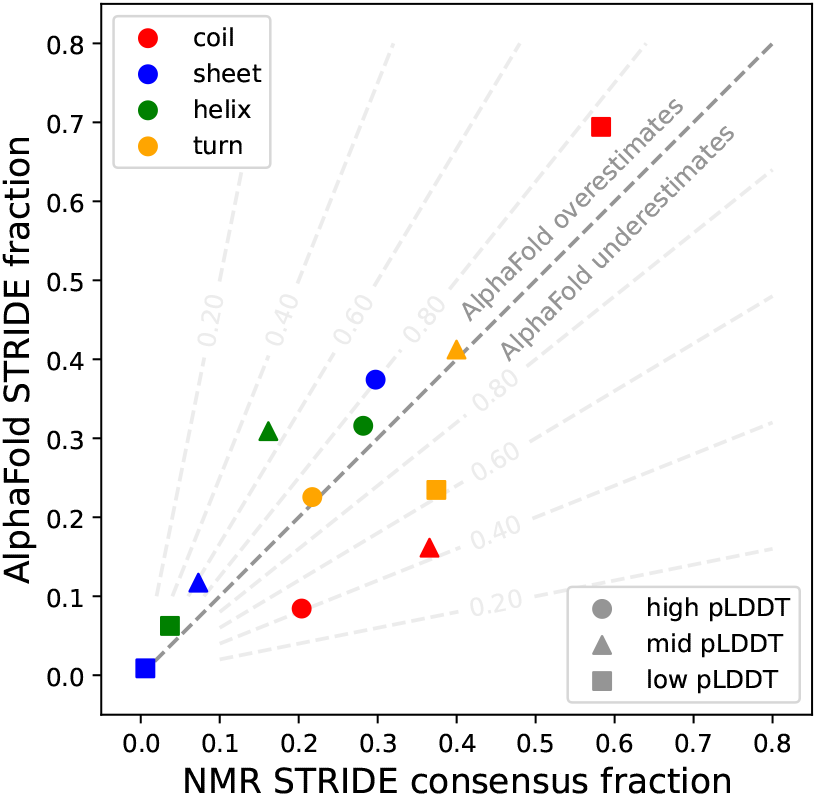
Comparison of STRIDE secondary structures fractions in AlphaFold2 models and NMR ensembles. Values are normalised to highlight the differences in secondary structure content (colour coded) captured per pLDDT class (coded by mark). Values above the dotted line indicate a higher presence in AlphaFold2 models, values below the dotted line indicate a higher presence in NMR ensembles. Based on 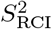 dataset.

For residues with low pLDDT values, which are not confidently predicted by AlphaFold2, a higher tendency for residues with coil secondary structure is present than observed in the NMR ensembles, with β-sheet and helix very similar in content and turn underrepresented in the AlphaFold2 models. This indicates that the AlphaFold2 models are not able to capture transient turn conformations that are still modelled in the NMR ensembles, where sufficient experimental data capturing distances and/or dihedral angles can be present to detect such conformations. For the mid pLDDT values, helix is overrepresented in AlphaFold2 models, whilst coil is underrepresented with respect to NMR ensembles. This discrepancy likely points towards regions that are in coil conformation but can fold upon binding to an interaction partner, evidenced by the observation of single isolated helices in many AlphaFold2 models [44]. Finally, for high pLDDT values, β-sheet is overrepresented and coil underrepresented in the AlphaFold2 models compared to NMR ensembles. A likely explanation here is that residue interactions in β-sheets are often between residues far removed in the sequence, with information from NMR experiments therefore often sparser and more likely to be insufficient to define such β-sheets with enough precision in the calculated structure models. Secondary structure classification programs such as STRIDE are then less likely to unambiguously assign them, even if they are present in solution.

Comparing the STRIDE secondary structure assignments of the 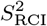 dataset at the residue level (Fig. 2) reveals that the discrepancies in secondary structure assignment increase as the pLDDT decreases. Notably, coil and turn residues often swap assignments with each other at low pLDDT. The low confidence in the AlphaFold2 predictions of these residues (supplementary Fig. 9) affects the interpretation of these assignment swaps. The presence or absence of a unique STRIDE assignment for the corresponding residue in an NMR ensemble is also important, with the unique assignments corresponding to higher pLDDT values (Supplementary Figs. 9, 10 and 11). Transient turn conformations that are present in NMR ensembles are often not captured by AlphaFold2, whereas AlphaFold2 predicts turns where the NMR ensembles do not show any (Fig. 2). Turn conformations are essentially locally determined, with often limited information available to precisely define them. Experimental NMR information can therefore be insufficient to precisely define turns in the NMR ensembles, while the required combination of co-evolutionary and encoded structure information might on the other hand not be present for a confident turn prediction in AlphaFold2 models.

**Figure 2:**
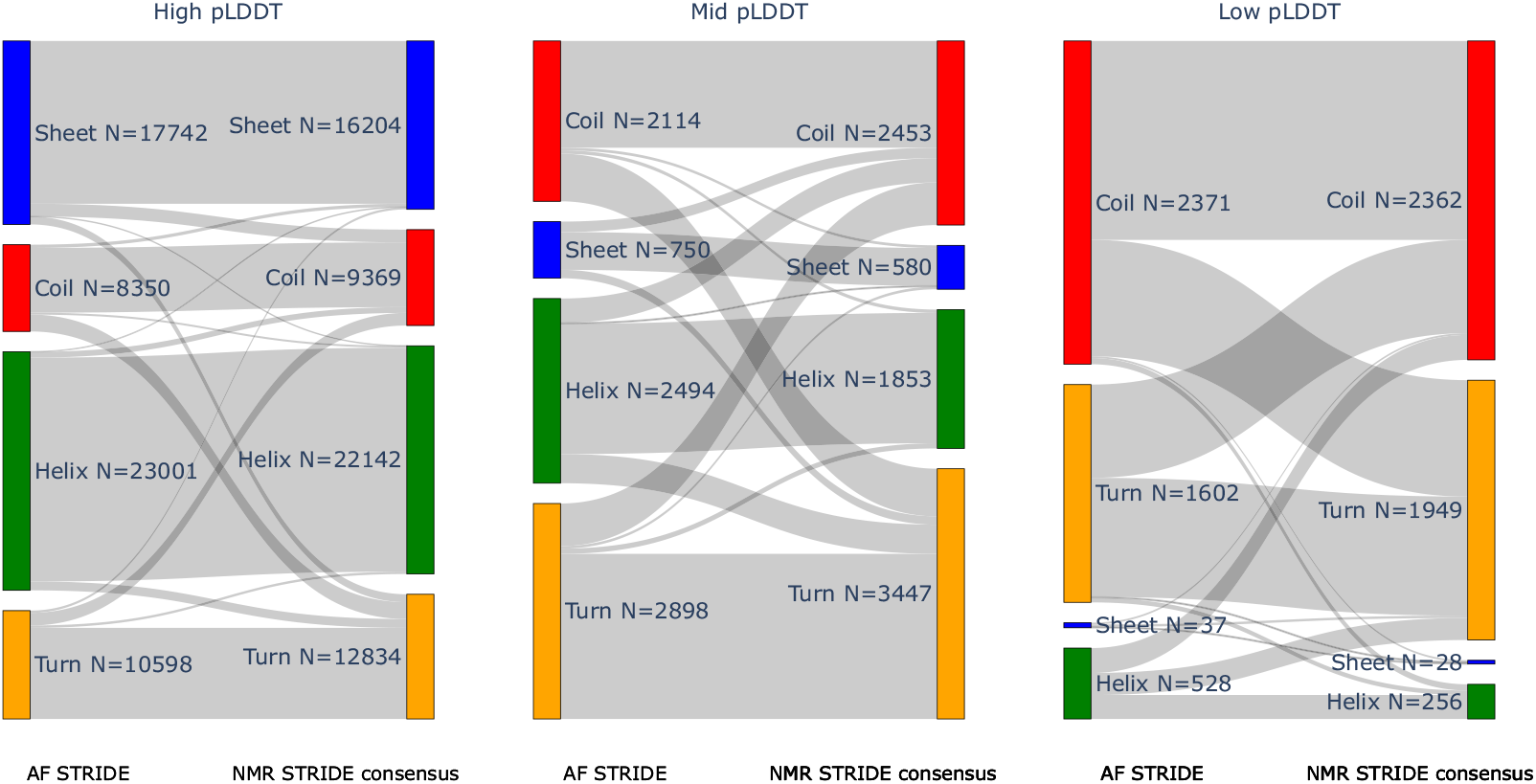
Per-residue comparison of STRIDE secondary structure assignment in AlphaFold2 models and NMR ensembles. For each pLDDT class (left, middle, right) the correspondence between the STRIDE secondary structure assignment for the AlphaFold2 (coloured bars on left side of each plot) and NMR ensembles (right side of each plot) is shown. Gray bars indicate the similarities and changes at the residue level between the STRIDE assignments, with no normalisation applied. Based on 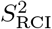 dataset.

### 3.2 AlphaFold2 pLDDT vs NMR experimental order parameters

Information on the dynamics of individual residues for proteins in solution, at near-physiological temperatures, can be directly captured by the NMR *S*^2^ order parameter. It can also be derived from the NMR chemical shifts of these residues, as done by the RCI method, which interprets chemical shifts to estimate the 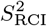 proxy for the order parameter. The RCI method uses a simple linear model to compare observed chemical shift values to reference ‘random coil’ chemical shift values, which indicate many highly dynamic conformations. We here compare these measurements to AlphaFold2 structures, which only offer a snapshot structure of a protein that in itself does not contain information about its motions. Given that AlphaFold2’s pLDDT metric is a good predictor for disorder [15], with a clear relationship between the pLDDT and amino acid order/disorder preference (Supplementary Fig. 1), we here explore to which extent this parameter also captures the degree of dynamics a residue experiences.

#### 3.2.1 Relation between chemical shift derived 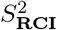 backbone dynamics and pLDDT values

The relationship between the pLDDT values and their corresponding RCI-derived order parameter 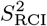 (Fig. 3 A & B) shows an unambiguous tendency for residues confidently predicted by AlphaFold2 (high pLDDT) to be also highly ordered in solution (high 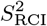, above 0.80). Conversely, residues not confidently predicted by AlphaFold2 (low pLDDT values) tend to be highly dynamic in solution (low 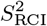, below 0.80) with a wide spread of 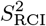 values. Mid pLDDT values display a hybrid behaviour, with a peak of higher 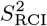 values complemented by a long tail of lower 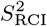 values that tailor off towards low 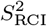. These results show that, as expected, the regions of proteins that are well folded and rigid in solution are accurately predicted by AlphaFold2, with the secondary structure of the experimental models in close agreement (see section 3.1). Residues predicted with medium pLDDT cover both highly rigid to highly flexible residues, reflecting an undefined relationship with in-solution protein behavior. Here, AlphaFold2 is not able to distinguish the extent of dynamics present, though it captures, to some degree, the associated conformational ambiguity of such residues by the generally lower pLDDT values. Finally, low pLDDT values unambiguously correspond to residues that are dynamic in solution. The Pearson correlation between pLDDT and 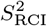 on the full dataset is relatively high (Table 2, top row), but this is driven by very high pLDD values with high 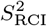 and very low pLDDT values with low 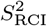. When considering each of the pLDDT categories, only very limited correlations are observed (Table 2, second grouped rows). This is even more pronounced when looking at 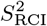 stratified classes, where almost no correlation remains (Table 2, fifth grouped rows). The 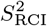 values are here subgrouped as flexible (below 0.70), rigid (above 0.80), and context-dependent ambiguous (0.70-0.80), as defined previously [45]. Overall, these results indicate that AlphaFold2 picks up on extensive conformational variability, but does not capture the amount of movement between such multiple conformations. Mann-Whitney two-sided U tests confirm significant differences between all subsets with a p-value *<* 0.001 (Supplementary Table 3).

**Table 2:** Overview of Pearson correlation for diverse metrics at different pLDDT and 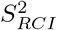 ranges. *Note: p-values marked with * were too low for Scipy to differentiate from 0*.

| Metric A | Metric B | Stratification Metric | Stratus range | Pearson's correlation | P-value | n |
| --- | --- | --- | --- | --- | --- | --- |
| pLDDT | $S^2_{RCI}$ | None | Unstratified | 0.6727 | 0* | 76,711 |
| pLDDT | RMSF | None | Unstratified | -0.5029 | 0* | 338,301 |
| $S^2_{RCI}$ | RMSF | None | Unstratified | -0.2197 | 0* | 73,814 |
| pLDDT | $S^2_{RCI}$ | pLDDT | Low | 0.3129 | $5.21 \times -119$ | 5,223 |
| pLDDT | $S^2_{RCI}$ | pLDDT | Mid | 0.3123 | $1.16 \times -199$ | 8,859 |
| pLDDT | $S^2_{RCI}$ | pLDDT | High | 0.2348 | 0* | 62,629 |
| pLDDT | RMSF | pLDDT | Low | -0.0359 | $1.93 \times -26$ | 87,690 |
| pLDDT | RMSF | pLDDT | Mid | -0.0721 | $2.99 \times -47$ | 40,041 |
| pLDDT | RMSF | pLDDT | High | -0.1272 | 0* | 210,570 |
| $S^2_{RCI}$ | RMSF | pLDDT | Low | -0.2001 | $1.89 \times -32$ | 3,445 |
| $S^2_{RCI}$ | RMSF | pLDDT | Mid | -0.1262 | $3.34 \times -30$ | 8,128 |
| $S^2_{RCI}$ | RMSF | pLDDT | High | -0.1262 | $3.34 \times -210$ | 62,241 |
| pLDDT | $S^2_{RCI}$ | $S^2_{RCI}$ | Flexible | 0.4772 | 0* | 13,656 |
| pLDDT | $S^2_{RCI}$ | $S^2_{RCI}$ | Context-dependent | 0.1547 | $2.26 \times -73$ | 13,560 |
| pLDDT | $S^2_{RCI}$ | $S^2_{RCI}$ | Rigid | 0.1841 | 0* | 49,495 |
| pLDDT | RMSF | $S^2_{RCI}$ | Flexible | -0.2348 | $5.39 \times -139$ | 11,113 |
| pLDDT | RMSF | $S^2_{RCI}$ | Context-dependent | -0.1118 | $2.80 \times -38$ | 13,314 |
| pLDDT | RMSF | $S^2_{RCI}$ | Rigid | -0.1402 | $2.56 \times -215$ | 49,387 |
| $S^2_{RCI}$ | RMSF | $S^2_{RCI}$ | Flexible | -0.1724 | $6.96 \times -75$ | 11,113 |
| $S^2_{RCI}$ | RMSF | $S^2_{RCI}$ | Context-dependent | -0.0542 | $4.03 \times -10$ | 13,314 |
| $S^2_{RCI}$ | RMSF | $S^2_{RCI}$ | Rigid | -0.0627 | $3.59 \times -44$ | 49,387 |

**Figure 3:**
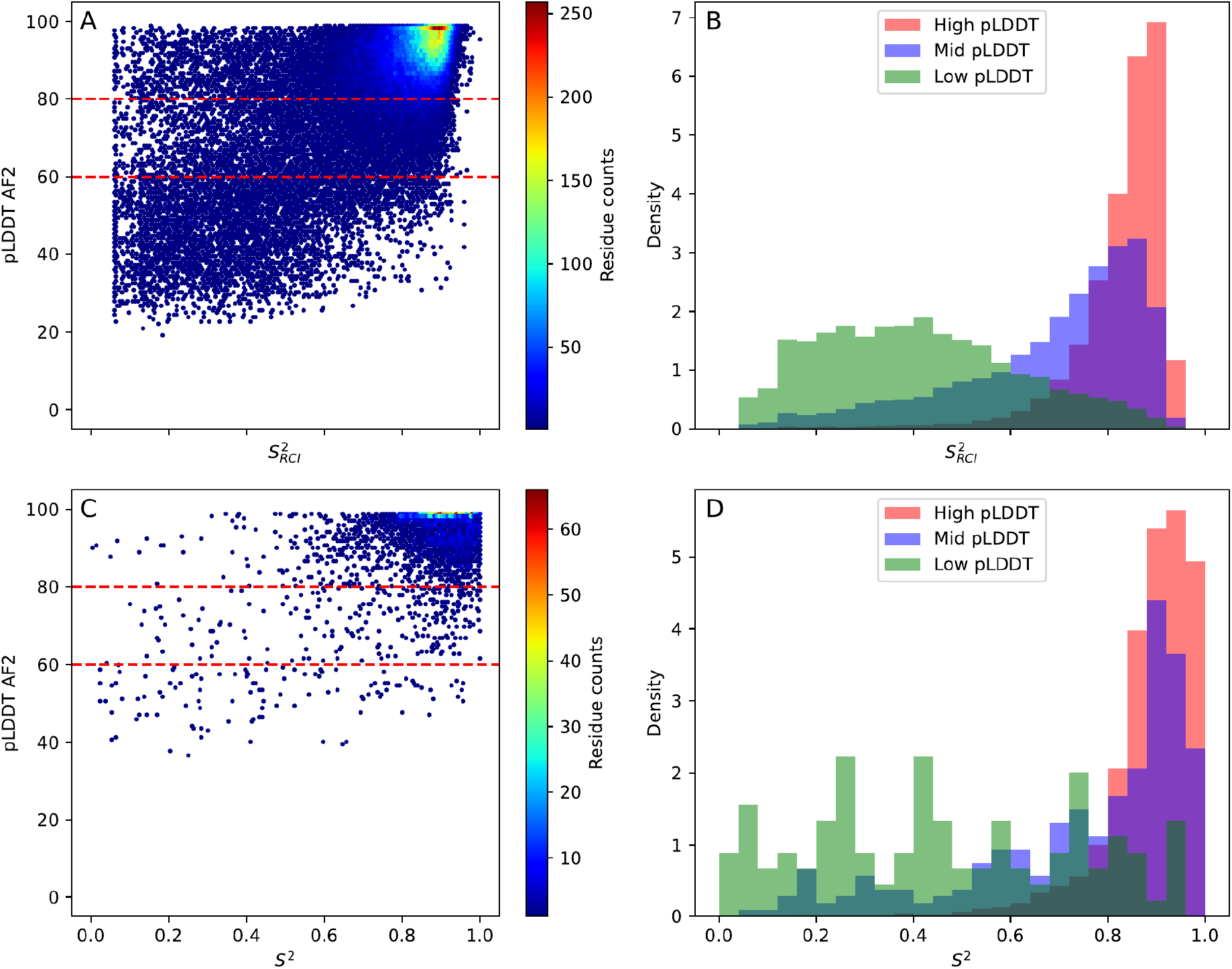
Comparison of pLDDT vs 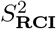 and *S*^2^. (A and C) Per-residue hexagonal binning plot of pLDDT versus order parameter values, with dotted red lines indicating cutoffs for classification of pLDDT values into three groups. The colour-coding indicates the residue count. Three groups for pLDDT 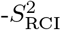 plot (A): high pLDDT (N=62,624), mid pLDDT (N=8,859), low pLDDT (N=5,223). Three groups for pLDDT-*S*^2^ plot (C): high pLDDT (N=4,103), mid pLDDT, (N=267), low pLDDT (N=112). (B and D) Histograms of order parameter value distributions for each pLDDT range.

#### 3.2.2 Relation between experimental *S*^2^ order parameter and pLDDT values

The 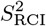 values are obtained from chemical shift data, which are readily available from the BMRB. The actual *S*^2^ order parameter, however, is much more difficult to measure with NMR, with as a result very few *S*^2^ values available from the BMRB. There are fundamental differences between the 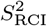 and *S*^2^ values, with the former less accurate and capturing movements on longer timescales (up to low ms), while the latter is highly accurate but only captures very fast movements (on the ps-ns timescale). Nevertheless, the relationship between the *S*^2^ order parameters and the pLDDT is similar to the one observed for the *S*^2^ (Pearson = 0.51, p-value 4.8*×*10^*−*289^), though the smaller dataset size results in noisier distributions (Fig. 3 C & D). The mid pLDDT value range is here more skewed towards *S*^2^ values higher than 0.8. This could be due to bias in this limited dataset, or could indicate that the ambiguous behavior described in section 3.2.1 is more relevant for slower (towards ms) movements, rather than very fast ones, as previously described in an analysis of 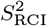 versus *S*^2^ values [35]. Moreover, very equivalent behaviour is observed for the more recent AlphaFold3’s C_α_ pLDDT (Supplementary Fig. 8). Although there are significant changes in the AlphaFold2AlphaFold3 pLDDT for individual residues within this dataset, the overall statistics do not change. Here also Mann-Whitney two-sided U tests confirm significant differences between all subsets with a p-value *<* 0.001 (Supplementary Table 3)

### 3.2.3 Relation between chemical-shift derived *δ*2D secondary structure populations and pLDDT values

The *δ*2D populations [46] are an estimation of the per-residue secondary structure occupancy, as fraction of 1.0, derived from NMR chemical shift data of the protein in solution. This method works similarly to the RCI method, except that *δ*2D interprets chemical shift values with a model that relies on chemical shift values typically observed for secondary structure elements. The available *δ*2D populations in the 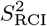 dataset were compared to the pLDDT values, similar to sections 3.2.1 and 3.2.2 (Supplementary Fig. 3 & Supplementary Table 2). The residues that according to *δ*2D adopt dominant helix and sheet populations are also typically predicted with high confidence by AlphaFold2. This is expected as both helix and sheet are rigid hydrogen bond stabilised secondary structures. Note that these secondary structures are mutually exclusive, with high helix population corresponding to low sheet population, which accounts for the high proportions of zero occupancy in Supplementary Fig. 3. In contrast, coil and polyproline II (PPII) populations, which feature multiple conformations and variable hydrogen bonding, are predicted with low confidence by AlphaFold2 (low pLDDT). These results are therefore in line with the previous observations from Fig. 2 & Supplementary Fig. 2.

#### 3.2.4 Relation between ShiftCrypt chemical shift interpretation and pLDDT values

ShiftCrypt [23] is a machine learning based method that encodes chemical shift values in values between 0 and 1, encompassing a combination of conformation and dynamics from rigid helix (towards 0.0) and rigid sheet (towards 1.0), with values around 0.5 indicating dynamic behavior and multiple conformations. The method was employed on the chemical shift data in the 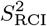 dataset and compared to pLDDT and stratified ranges (Fig. 4 panels A & B). High pLDDT values are associated with high and low ShiftCrypt values (indicating rigid sheet and helix), while low pLDDT ranges are almost exclusively in the 0.4-0.6 ShiftCrypt range (indicating multiple conformations in dynamic exchange). These results therefore confirm those of the other chemical shift based methods previously discussed.

**Figure 4:**
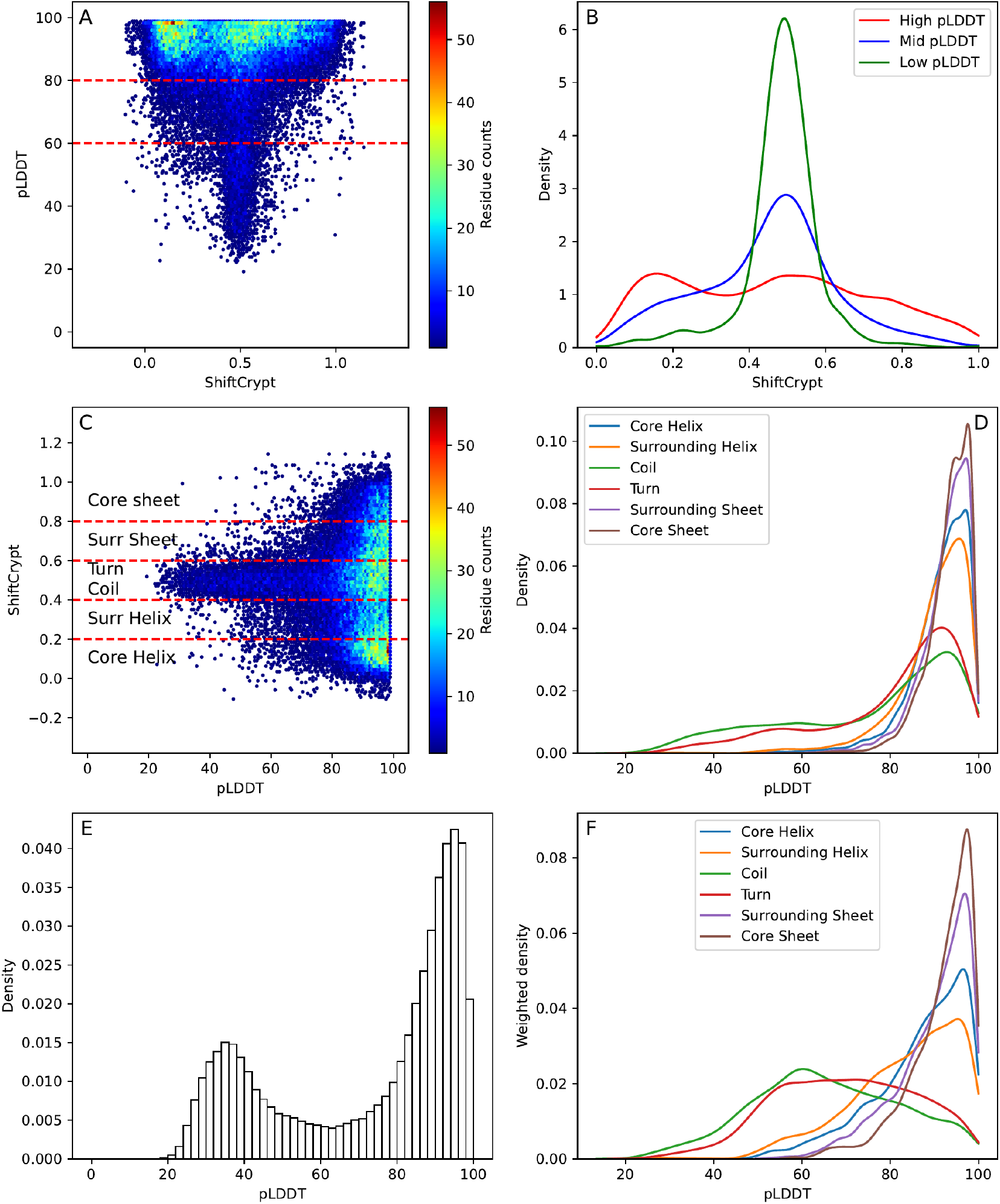
Analysis of pLDDT and ShiftCrypt values. A & B: The residues with valid ShiftCrypt values were compared with their corresponding pLDDT and stratified according to their pLDDT values in high (N=41,293), mid (N=5,480) and low (N=2,938). C & D: The residues of the comparison in panels A and B were divided with the same criteria to determine the Constava conformational states [36]: core helix (N=8,122), surrounding helix (N=5,698), coil (N=5,236), turn (N=5,854), surrounding sheet (N=4,947), core sheet (N=3,563). E & F: The distributions in panel D were weighted by the inverse of the overall pLDDT density in the dataset, resulting in a weighted density of pLDDT values for each conformational state.

Conversely, the ShiftCrypt values combined with the STRIDE [39, 40] secondary structure assignment of the NMR ensembles can be used to categorise residues into 6 conformational states, as described by Gavalda-Garcia *et al*. [36]: core helix, core sheet, surrounding helix, surrounding sheet, coil, and turn. These conformational states comprise a combined interpretation of dynamics and conformation, with the “core” states highly rigid, the “surrounding” states more dynamic, and the “coil” and “turn” states highly dynamic. When comparing the conformational state of a residue with the pLDDT value of the residue (Fig. 4 C & D), it is again clear that low pLDDT values correspond to the turn and especially coil states, with likely multiple conformations in dynamic exchange present. Due to the skew of the overall distribution of pLDDT towards high values (Fig. 4 E), high pLDDT values dominate for all conformational states, though only the “coil” and “turn” states feature significant densities below pLDDT of 80%.

To address the skew, the densities of each conformational state were normalised by the total pLDDT density over all states to highlight relative pLDDT tendencies for each conformational state (Fig. 4 F). Both sheet states feature the largest relative densities of high pLDDT values, with few values lower than 80 and very few lower than 60, indicating that if they are present in solution, then they are confidently predicted by AlphaFold2. Since these conformations feature non-local structural stabilisation by hydrogen bonds with other *β*-strands, this restricts their motions and fixes their position, as well as leading to strong co-evolutionary signals [47–49], so facilitating structural predictions. Helix conformations follow sheet in high pLDDT density, though their density is shifted towards lower pLDDT values. Helix conformations are stabilised by a regular hydrogen bond network defined by local interactions between amino acids, making them easier to predict and design than sheet [50]. The strong local sequence dependence implies that even when helices are more dynamic and partially unfold, they are easier to reform [51–53], with co-evolutionary signals also harder to pick up [54]. Both of these factors can here explain the observed difference with sheet. Overall, the “core” helix and sheet correspond to higher pLDDT than the “surrounding” states, again highlighting that the pLDDT does in this case pick up the likelihood of multiple conformations being present. Finally, the coil and turn states are quite evenly spread over the whole pLDDT range, indicating that these residues, which are likely to adopt multiple conformations, can still be confidently predicted, either due to stringent evolutionary constraints or due to single defined conformations being observed in the AlphaFold2 training set.

An example of where AlphaFold2 might confidently predict structure, but the ShiftCrypt values indicate multiple conformations, are protein regions that fold upon binding. Indeed, the transformer-based model architecture and large dataset employed to train AlphaFold2 captures intrinsic properties of protein folds and the role of amino acids, such as a residue’s location (*e*.*g*. hydrophobic amino acids would favor protein core location), which consequently affect their degrees of freedom and conformation. Since in the pre-processing of AlphaFold2’s training dataset, all binders were removed to obtain a monomer-only set prior to training, this has significant consequences for regions of proteins that fold upon binding. Such regions only acquire a fold in their bound form [55], which is then what AlphaFold2 tends to predict [56], so enabling the placement of for example co-factors in AlphaFold2predicted structures [57]. These regions should show conflicting, more dynamic behaviours than those expected from pLDDT, as captured in NMR-derived metrics from the unbound proteins. Indeed, Fig. 4 shows that there are a considerable number of residues with high pLDDT and mid ShiftCrypt values, which indicate dynamic resides that were easy to predict by AlphaFold2.

To investigate these occurrences, all proteins were scanned for stretches of at least 15 consecutive residues with pLLDT >80 and 0.4 <ShiftCrypt <0.6. As example, we selected the mouse Cytohesin3 or Grp1 (Uniprot accession code O08967), which features self-inhibition in the absence of a ligand, and is only active in the presence of Ins(1,3,4,5)P_4_ or an analogous molecule [58]. Fig. 5 A shows the predicted AlphaFold2 structure with a loop featuring high pLDDT that is unstructured according to ShiftCrypt. Fig. 5 B shows the experimentally determined structure, which illustrates that this loop is located where Ins(1,3,4,5)P_4_ binds the PH domain if Grp1. All publicly available experimental structures of this protein or domain include Ins(1,3,4,5)P_4_ or a fragment of it in their structure (PDB IDs: 1FGY, 1FGZ, 1FHW, 1FHX, 1U2B, 2R09, 2R0D, 6BBP & 6BBQ), all published before April 2018 and so part of AlphaFold2’s training set. In contrast, the ShiftCrypt value was calculated from chemical shift data (BMRB ID 15669), which did not include any ligand, with the loop unbound and accessible to solvent (Supplementary Fig. 5). The dynamical behavior is confirmed by low 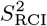 (data in supplementary data frames https://zenodo.org/doi/10.5281/zenodo.10977724) and highlights the limitations of even highly-confident predictions to capture ambiguous, context-dependent behavior of protein regions.

**Figure 5:**
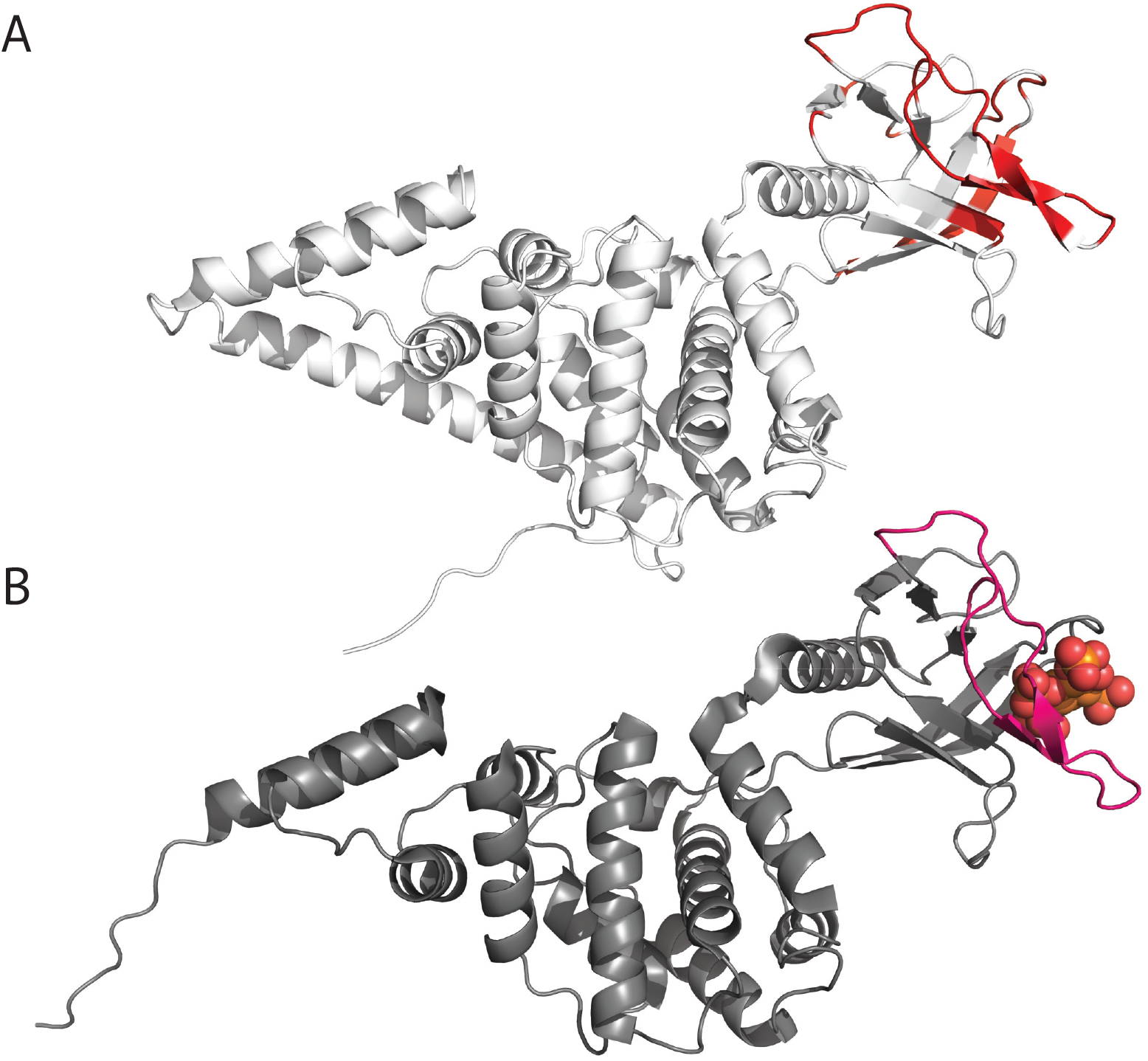
Example of long region with conflicting pLDDT and ShiftCrypt values. Above: High pLDDT with mid ShiftCrypt values. Below: Experimental structure (PDB: 2R0D) with Ins(1,3,4,5)P_4_ bound. Highlighted is the loop which features a long continuous segment of high pLDDT with mid ShiftCrypt values. All other experimental structures available in PDB also featured the presence of Ins(1,3,4,5)P_4_ or a fragment of it (not shown).

### 3.3 Low pLDDT values are almost exclusively found in regions with high conformational state variability in molecular dynamics trajectories

The MD dataset, comprising MD trajectories for 100 proteins, was analyzed using the conformational state variability metric, which measures the potential for a residue to exist in multiple conformational states. Each conformational state represents a probabilistically defined macro-state that a residue can adopt, defined from NMR data as probability density functions in dihedral space, and represents different low energy conformational states that residues tend to adopt in solution [36]. Given a set of structures, such as from MD trajectories, the propensity for each conformational state (*i*.*e*. the degree of preference to adopt a conformational state) can be calculated for each residue, with the conformational state variability then indicating how likely it is for a residue to adopt multiple conformational states. For example, a fully disordered residue which samples the coil dihedral probabilities during an MD simulation could feature near-0 conformational state variability, as exclusively adopts a highly dynamic coil state. If it would intermittently adopt helix conformation and switch back to coil, its conformational state variability would in contrast be higher.

The residues of all proteins in the MD dataset (Table 1) were divided in pLDDT ranges and the distributions of the conformational state variability for each range were compared (Fig. 6 & Supplementary Fig. 4). This shows that residues with high pLDDT have low conformational state variability. Residues with low pLDDT have almost exclusively to high conformational state variability, confirming that such residues generally have the ability to exist in diverse conformational states. Mann-Whitney two-sided U test yielded a p-value *<* 0.001 between all distributions (Supplementary Table 3). The relatively small size of the dataset enabled us to confirm this relationship for the C_α_ pLDDT of AlphaFold3 models (Supplementary Figs. 6 & 7 & Supplementary Table 1).

**Figure 6:**
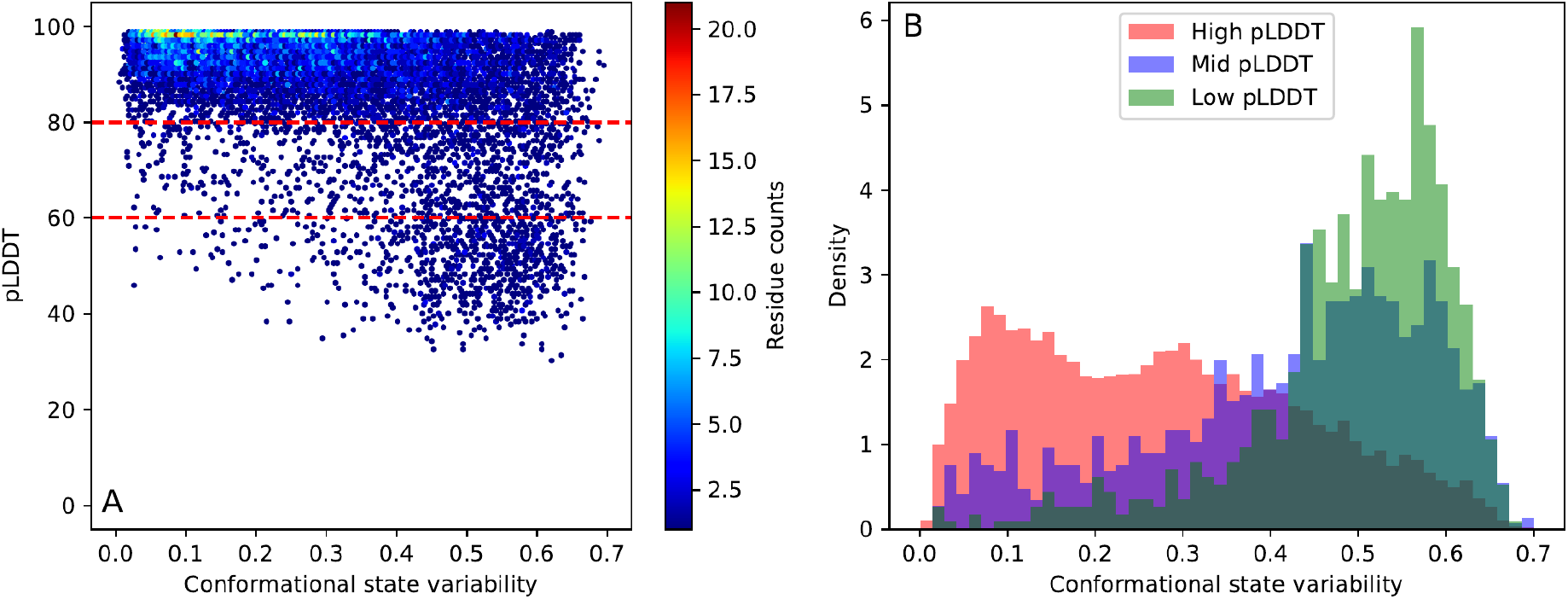
pLDDT vs conformational state variability. A) Per-residue hexagonal binning of pLDDT versus the conformational state variability, with high (N=9,523), medium (N=1,038) and low pLDDT (N=809) pLDDT regions indicated by red dotted lines. B) pLDDT-stratified distributions for each of these classes. The distributions for each conformational state propensity can be found in Supplementary Fig. 4 & Supplementary Table 1.

### 3.4 AlphaFold2 pLDDT vs NMA fluctuations of AlphaFold2 models

Another per-residue metric for protein flexibility are the root mean square fluctuations (RMSF). Using 200 normal modes from the NMA as obtained with the WEBnma tool, the RMSF were computed for the AlphaFold2 structures in the 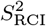 dataset based on the C_α_ positions (see Methods 2.1.2). We explored how well the pLDDT of the AlphaFold2 model captures fluctuations computed from NMA.

Using the standard NMA, as shown in Supplementary Fig. 12, some coil residues may exhibit extremely high RMSF reaching up to 185 Å, primarily in flexible Nand/or C-termini (Supplementary Table 4). Such high RMSF values can even occur in termini that are rigid units themselves. Indeed, when another segment of the protein experiences significant motion, this motion can propagate throughout the entire protein, even if those regions are densely packed and conformationally stable [26]. Thus, the presence of loose segments with substantial RMSF can shift the RMSF values for the entire protein to higher values. We therefore truncated loose termini and reran the NMA and RMSF calculations (see Methods 2.1.2). In comparison with the non-truncated models, these truncated AlphaFold2 3D structures had considerably fewer unphysical RMSF values (Supplementary Figs. 12 and 13). The further analysis in this paper pertains to the NMA fluctuations obtained from truncated AlphaFold2 models.

To compare the influence of secondary structure elements, the RMSF of the residues were grouped according to their STRIDE assignment, i.e. coil, strand, *α*-helix, turn, 3_10_-helix, or bridge. The statistics of the RMSF values (minimum, maximum, mean, and standard deviation (std)) are reported (Supplementary Table 5) for the high, mid, and low pLDDT residues. The two-dimensional histogram of the RMSF and pLDDT is visualized for coil and *α*-helix residues in Fig. 7A and 7B, respectively. For each secondary structure element, the Pearson correlation coefficient between RMSF and corresponding pLDDT residues was computed (Supplementary Table 7) to investigate their relationship. The reported correlations for pLDDT in Supplementary Table 7 are significant (p-value < 0.05) except for strand residues with mid pLDDT, 3_10_-helix residues with mid pLDDT, and bridge residues in low and mid pLDDT regions.

**Figure 7:**
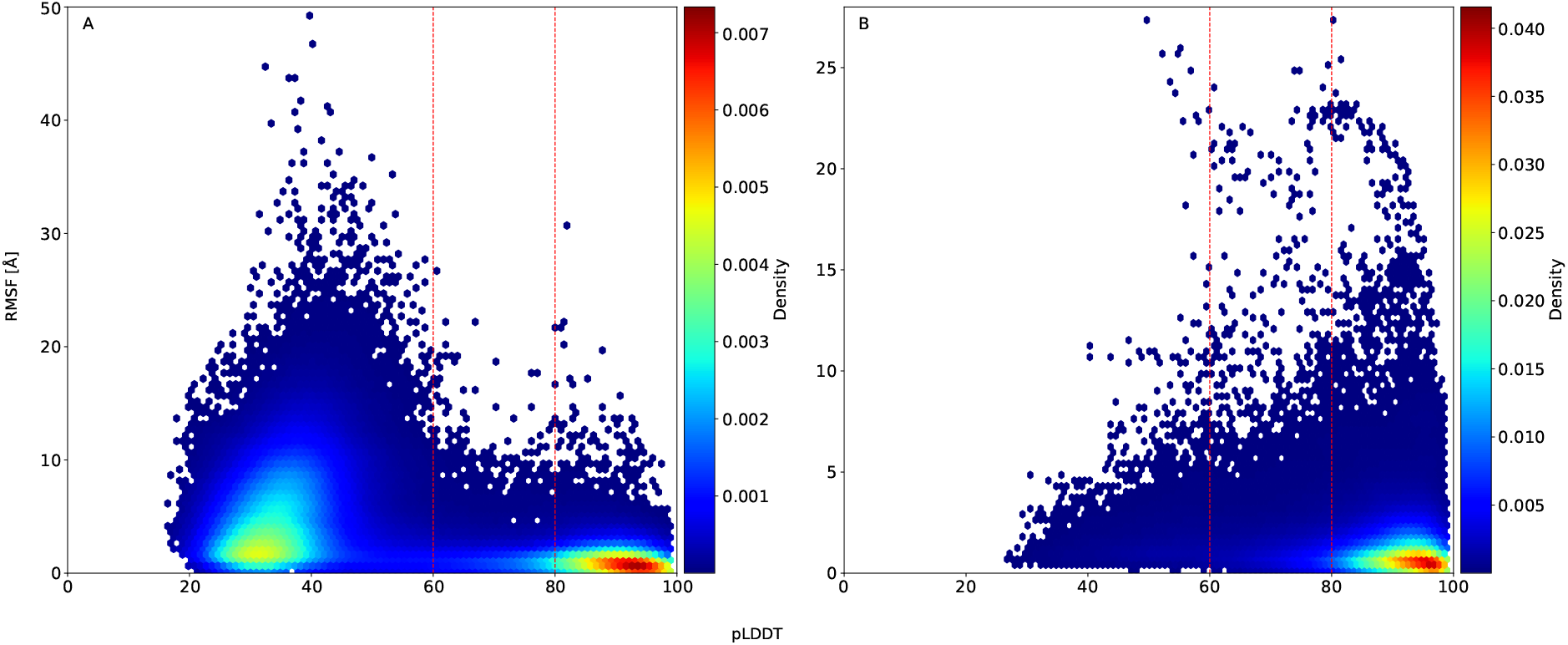
pLDDT vs RMSF of AlphaFold2 models. Histogram of pLDDT vs NMA fluctuations of each amino acid, visualized with a Gaussian kernel estimator. Data from 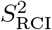 dataset. Subplots shown for secondary structures A) coil (N=105,172), B) *α*-helix (N=109,639), where N represents number of amino acid residues. The red vertical lines divide the dataset into high, mid, and low pLDDT regions.

For residues with low pLDDT, coil residues show highest mean RMSF of 5.65 *±* 4.43 Å compared to other secondary structure elements, indicating that coils have the highest flexibility of all secondary structures, as expected [59]. In contrast, the mean RMSF values for the other secondary structures vary, ranging from 1.89 Å for strands and bridges to 2.16 Å for turns (Supplementary Table 5, Supplementary Fig. 16).

For residues with mid pLDDT, coil, *α*-helix, and 3_10_-helix residues exhibit a slightly higher mean RMSF of 1.84 − 1.89 Å compared to the other secondary structure elements which fall within a similar range, ranging from 1.34 Å to 1.74 Å (Supplementary Table 5). The difference in mean RMSF of coil residues compared to other secondary structure elements is here minimal. Remarkably, mid pLDDT residues have a reduced mean RMSF compared to low pLDDT residues, independent of the secondary structure element. This points to a reduced flexibility for mid pLDDT residues compared to low pLDDT, which is in agreement with expectations.

Lastly, for residues with high pLDDT, *α*-helix residues show the highest mean RMSF of 1.38 Å, closely followed by the other secondary structure elements led by coil in the range 1.11-1.33 Å (Supplementary Table 5). Moreover, the mean RMSF is lowest for high pLDDT residues, in line with their higher expected rigidity.

Thus, in conclusion, coil residues are most flexible when they have a low AlphaFold2 pLDDT value, whereas coil and *α*-helical residues are most flexible when they have a higher pLDDT. Most interestingly, there is a consistent reduction in mean RMSF when the pLDDT increases, irrespective of the chosen secondary structural element. A weak negative correlation can indeed be found between RMSF and pLDDT with Pearson correlation coefficients ranging from − 0.11 (p-value=4.49 × 10^*−*319^) for *α*-helix residues to − 0.43 (p-value ≈ 0) for coil residues (Supplementary Table 7). Overall, the Pearson correlation coefficient between RMSF and pLDDT of all 338,301 residues, irrespective of secondary structure, is − 0.50 (p-value ≈ 0) (Table 2). The negative correlation can be understood by looking at the elastic network model (ENM) assumptions. The ENM models the intramolecular interactions as springs between C_α_ atoms, and therefore lower RMSF will correspond in general to regions with a higher spatial density of C_α_ atoms. This is in line with the observation that high pLDDT values are related to well-folded regions of a protein, with residues tightly packed [1].

However, this correlation is again driven by most pLDDT values being either high (>80) or low (<40), with correspondingly lower and higher RMSF values. While there is an weak overall negative Pearson correlation, this is not present when considering the different pLDDT or 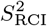 subgroups (Table 2, third and sixth grouped rows). Within secondary structure elements this lack of correlation holds. For example for coils, the mid pLDDT (−0.17, p-value 8.98 × 10^*−*55^) and high pLDDT (− 0.16, p-value 1.90 × 10^*−*157^) correlations are slightly negative (Supplementary Table 7), but for low pLDDT regions there is even a weak positive correlation (0.16, p-value ≈ 0.00). Upon visual examination of structures exhibiting these positive correlations, we mostly identified large proteins consisting of long, extended disordered coils connected by small segments of ordered regions. One possible explanation is that such regions might exhibit higher pLDDT than expected, compared to fully disordered regions, through their intermittent association with short folded regions with high pLDDT. Hence, while pLDDT is in general a reliable indicator of the RMSF (with overall Pearson correlation coefficient −0.50 with p-value ≈ 0), this is driven by the difference between very high and low pLDDT values and does not capture the intermediate flexibility gradations within each subgroups of low, mid, or high pLDDT.

### 3.5 NMA fluctuations vs 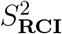

The relationship between the RMSF of the 3D structure models and the experimental 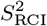 values, which give approximations of the backbone dynamics at the residue level, was investigated for the 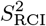 dataset (Supplementary Table 6, Supplementary Fig. 17). Residues were grouped according to their STRIDE secondary structure assignment. As for the 3D structures, each protein in the dataset had one AlphaFold2 model and a set of one or more NMR models (the NMR ensemble of the protein entry). If the NMA fluctuations on the model efficiently generate low-frequency normal modes capable of capturing protein motions commonly observed in NMR experiments, the NMA RMSF should negatively correlate with 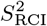 [28, 30], as higher values of RMSF indicate higher flexibility while higher values of 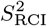 (close to 1) indicate rigid residues.

#### 3.5.1 Relationship of 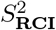 with NMA fluctuations on AlphaFold2 models

The overall Pearson correlation between RMSF and 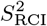 is indeed negative but very weak (Table 2), as illustrated for coil and *α*-helix residues (Figure 8). The negative correlations are less weak when considering only coil residues (Pearson = − 0.32, p-value = 1.06 × 10^*−*278^, see also Supplementary Table 7). This observation indicates that NMA on AlphaFold2 models does not capture the experimental 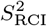 trends very well. The NMA fluctuations of the AlphaFold2 models are expected to be dominantly determined by the packing of the protein, which determines the structural flexibility to some extent. However, the 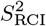 values indicate that this is not sufficient to capture the behavior of the protein in solution at higher temperatures. In fact, the pLDDT values are better correlated than the NMA RMSF (Table 2).

**Figure 8:**
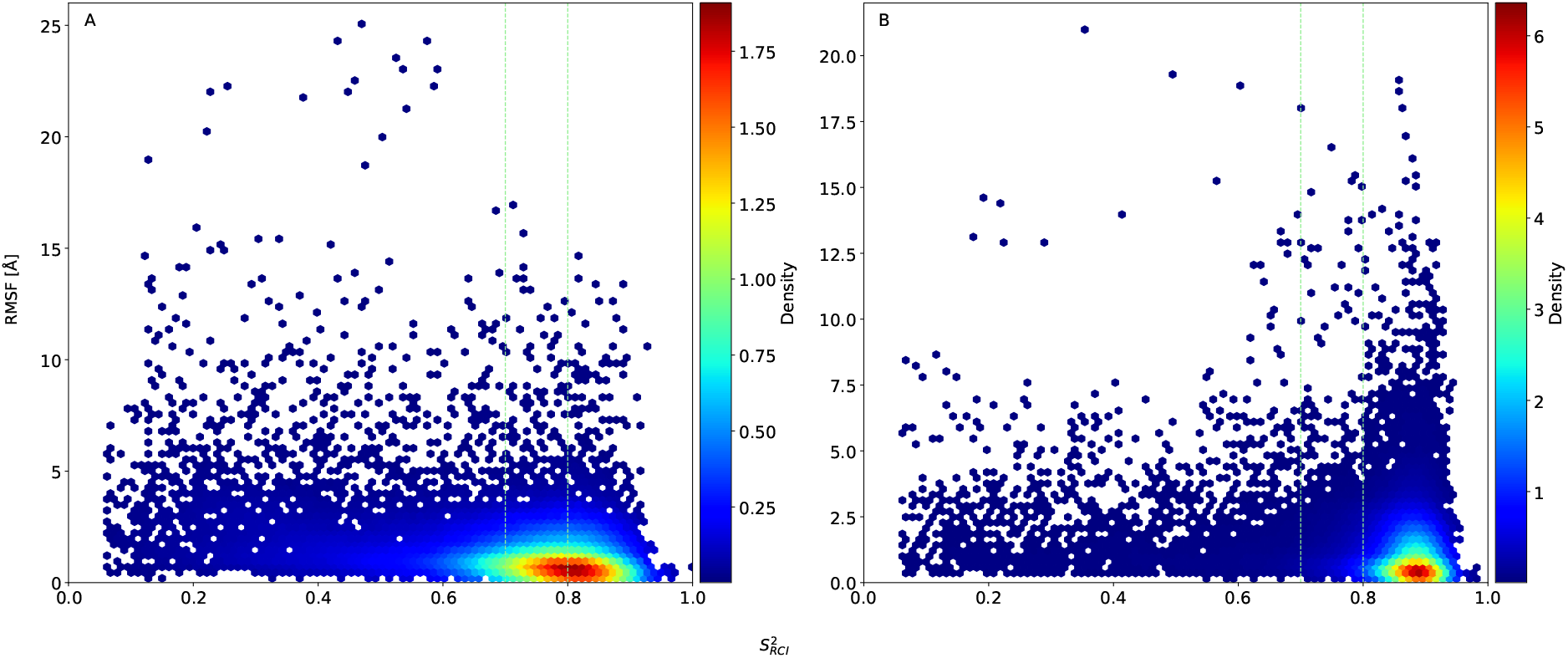
RMSF vs 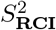. RMSF values versus 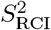 value of each amino acid, visualized with a Gaussian kernel estimator for truncated 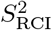 dataset. One subplot for each secondary structure element: A) coil (N=11,634), B) *α*-helix (N=25,861), where N represents number of amino acid residues. The green vertical lines divide the dataset into flexible 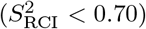, ambiguous 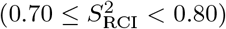, and rigid 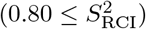 regions.

#### 3.5.2 Relationship of *S*^2^ with NMA fluctuations on NMR models

Every protein entry in the 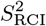 dataset not only has an AlphaFold2 model but also a set of one or more NMR models. The elastic network model was also built for each of the NMR models, and the NMA fluctuations were computed with WEBnma (Methods 2.1.2). The question arises whether the 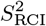 order parameter correlates significantly with the RMSF of the NMR models as well, despite the several differences between the AlphaFold2 and NMR models. AlphaFold2 structures are based on a machine learning prediction, trained on mostly crystallographic data at lower temperatures, while an NMR structure ensemble is typically generated by successive simulated annealing molecular dynamics simulations using a force field that incorporates restraints based on experimental NMR information for the protein in solution. The AlphaFold2 is therefore a single prediction, whereas the NMR ensemble is a set of structures that each encompass the experimental NMR information as best as possible. The NMR ensemble of a protein thus likely comprises more of the dynamics between the different protein conformational states.

To compare the NMR models with their corresponding AlphaFold2 models, the following procedure was executed. First, the Pearson correlation coefficient 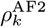 is computed between the 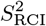 and the RMSF of its AlphaFold2 model, for each protein *k* individually (*k* = 1, …, 746). Second, for the same protein *k*, the Pearson correlation coefficient 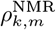 is computed between the 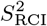 and the RMSF of each model *m* in its NMR ensemble. Hence, for a given protein *k*, this yields one correlation 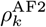 and a set of one or more correlations 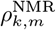. For the total 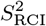 dataset, we obtained 746 Pearson correlation coefficients for AlphFold2 models and 14,069 for individual NMR models (distribution of coefficients visualized in Supplementary Fig. 18). Each NMR coefficient 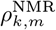 is paired with the corresponding AlphaFold2 coefficient 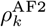, repeating the AlphaFold2 value when the NMR ensemble contains more than one model. The procedure thus results in 14,069 AlphaFold2-NMR pairs of models and their Pearson correlation coefficient between RMSF and 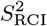 order parameters.

To determine whether 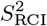 correlates differently with the NMA fluctuations of AlphaFold2 models or of the NMR models, a non-parametric Wilcoxon signed-rank test was performed on the distributions of these correlation coefficients. With a p-value of 0.0001, it indicates a significant difference between the AlphaFold2 and NMR model correlation coefficients, with the NMA fluctuations of the NMR models exhibiting a stronger negative correlation with 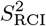 than the AlphaFold2 models. Specifically, among these pairs, 71.22% exhibited a stronger negative correlation for the NMR models compared to the AlphaFold2 model, and only 28.78% showed a weaker negative correlation. This indicates that NMA can better capture the actual dynamics of proteins in solution with multiple input models that better reflect the uncertainty in conformation, compared to NMA on a single AlphaFold2 structure model.

The different correlation coefficients must derive from a difference in the 3D geometry of these models, as the RMSF follows from an elastic network model attached to the C_α_ positions in the protein’s structure, and thus changes in RMSF originate from changes in packing and organization of the protein. We therefore compared the secondary structure assignments of the AlphaFold2 model and the NMR models using the STRIDE secondary structure assignments of each residue in an overlapping region of the AlphaFold2 sequence and NMR model sequence. Three situations can occur for a residue: the assignments are fully identical, fully conflicting, or ambiguous (given that multiple models are present in each NMR ensemble) (Supplementary Fig. 24). The percentage of conflicting residues in AlphaFold2-NMR pairs is then calculated as the ratio between the number of conflicting STRIDE assignments and the total number of residues that overlap between AlphaFold2 and NMR models. The percentage of conflicting STRIDE residues ranges from 0.86% to 88% of the total overlapping sequence length for 14,006 out of 14,069 pairs. However, the number of conflicting residues did not influence the correlation between 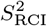 and RMSF (Supplementary Fig. 26). In addition, the structural size difference between AlphaFold2 and NMR models did not impact the flexibility profiles computed from WEBnma. This conclusion is supported by data from several proteins (Supplementary Table 8), which, despite having a few hundred more residues than the NMR structures, exhibited similar or even stronger negative correlations between RMSF and 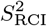 than NMR.

Overall, our findings suggest that conflicting secondary structure assignments between AlphaFold2 and NMR models, which indicate ambiguity in the behavior of that residue, do not directly impact the correlation between 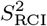 and RMSF (Supplementary Fig. 25). However, for individual proteins, conflicting residues within functionally relevant regions may significantly influence RMSF, especially in flexible regions. In instances where AlphaFold2 models show a positive correlation between 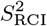 and RMSF, unlike the negative correlation seen in NMR models, conflicts in secondary structure predictions significantly impact this relationship. This discrepancy often occurs in regions where NMR identifies coils and turns while AlphaFold2 predicts different secondary structures. Interestingly, AlphaFold2’s high confidence levels do not mitigate the affect of these conflicts on RMSF; regions with high, mid, and low pLDDT scores exhibit similar impacts on RMSF and correlation (Supplementary Fig. 21).

An example illustrating this diversity is P0AFW0-2LCL, which showcases significant secondary structure differences between the NMR and AlphaFold2 models (88% of its overlapping AlphaFold2NMR sequence residues) (Supplementary Fig. 23). The Pearson correlation coefficients are − 0.55 for the AlphaFold2 model and − 0.74 to − 0.86 for the NMR models, with the conflicting residues occurring in low, mid, and high pLDDT regions. Another example is Q922K9-2D8J, where AlphaFold2 showed a stronger negative correlation than the NMR models despite conflicting secondary structure (Supplementary Fig. 20), with closer examination confirming that the effect varies from one protein to another (Supplementary Figs. 19, 21, and 22), with no discernible general trend based solely on secondary structures. Instead, the performance of AlphaFold2 and NMR models in capturing the relationship between 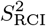 and RMSF appears to be very dependent on the specific characteristics and localization of conflicting residues within each protein.

In conclusion, in some instances, AlphaFold2 models seem to outperform NMR models, while in majority, NMR models outperform AlphaFold2 models. These results emphasize the importance of 1) validating the potential secondary structure conflicts between AlphaFold2 and NMR models, even in regions where AlphaFold2 shows high confidence, and 2) leveraging the relationship between 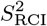 and RMSF to serve as a guide for refining these conflicts and improving the accuracy of AlphaFold2 models in capturing true solution dynamics.

### 3.6 Interactive analysis of entries

The entries for all datasets described in this work were processed to build a per-protein interactive analysis and made available on https://bio2byte.be/af_nmr_nma/. This resource offers a focused analysis of our entries, notably offering a dynamic mapping of the 3D structures and a selection of biophysical metrics, plotted along the amino acid sequence. Both the structure as well as the source data for any given protein can also be downloaded from their entry view, allowing users to visualize their other biophysical metrics, or to run further analysis on the datasets.

## 4 Discussion

This work presents a large-scale study of whether the AlphaFold2 pLDDT metric and models, through NMA analysis, can capture protein dynamics. The overall results confirm previous studies that, in general, residues with high pLDDT values are situated in well-folded, stable, rigid regions of proteins, whereas residues with low pLDDT are likely disordered and highly dynamic [15, 60, 61]. However, our results give a more complex and nuanced view of this relationship, linked to two key notions at the amino acid residue level that are dependent on the full protein. The first notion is “how many distinct conformations can a residue adopt?”, which is primarily thermodynamically determined. This depends on the low energy regions present in the complex energy landscape of the full protein, as well as on how this landscape changes with the overall conformation and the environment of that protein (e.g. folding upon binding due to the presence of a binding partner). The second notion is “how dynamic is a residue?”, which is kinetically determined, in other words by the height of the energy barriers between the low energy conformations. This will determine *how often* a residue can move between multiple accessible low energy conformations. Although these concepts are tightly linked, they are distinct from each other; a residue that adopts two distinct low energy conformations could move between them very quickly, very slowly, or anything in between, depending on the energy barrier between them. The observations from NMR that we compare to give the macroscopic view of such behavior, as an average of up to low ms timescales movements over the billions of molecular copies of the protein in solution, each with their individual behavior.

With this distinction in mind, the conflicts between the secondary structure of residues as observed in AlphaFold2 models versus NMR ensembles (Figs. 1 and 2) show that residues predicted with high and mid pLDDT as helix and sheet are often designated as coil in the NMR ensemble, whereas low pLDDT residues that are in turn conformation in the NMR ensembles are often predicted as coil by AlphaFold2. Interestingly, as evident from Supplementary Fig. 9 A/D/G, residues that have conflicts in secondary structure assignment tend to have lower pLDDT values. If we assume that such conflicts indicate that such a residue has a higher likelihood of adopting multiple conformations, these results show that AlphaFold2 detects the possibility of multiple conformations being present through lower pLDDT values. It is also evident from B/E/H and C/F/I in Supplementary Fig. 9 that these conflicting residues tend to be more dynamic in solution, with lower 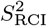 values and ShiftCrypt values between 0.4 and 0.6.

Another indicator of the presence of multiple conformations is when a residue has multiple secondary structure states over the structure models that compose an NMR ensemble. This indicates that the experimental NMR data used in the structure calculation was incompatible and/or insufficient to uniquely define the residue’s conformation. The overall trends are here similar to the AlphaFold2 conflicting residues, with the residues that do not have a unique secondary structure state throughout the NMR ensemble having lower pLDDT and 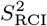 values as well as a large proportion of ShiftCrypt values between 0.4-0.6 (Supplementary Fig. 11). When in addition distinguishing by AlphaFold2 conflicting residues, a more complex picture emerges. For helix and sheet, unique NMR ensemble secondary structure assignments that match the AlphaFold2 model assignment are, according to in solution observations, solidly in the rigid helix (values towards 0) or sheet (values towards 1) categories regions (Supplementary Fig. 9). Even when the secondary structure information from the NMR ensemble is inconsistent, a matching AlphaFold2 assignment with the consensus NMR assignment still tends to reliably capture rigid helix and sheet present in solution. This might explain the observation that AlphaFold2 models often better encompass experimental NMR information than the calculated NMR ensembles themselves [62], as secondary structure elements will be well defined in AlphaFold2 models compared to NMR ensembles, for which the experimental NMR data might be insufficient to accurately form those secondary structures. Conversely, mismatches between the NMR ensemble and AlphaFold2 assignments for helix and/or sheet assignments are strongly indicative of dynamic behavior, with multiple conformations in solution. Note that our analysis attempts to highlight the differences with AlphaFold2 based on simple observations from NMR ensembles and experimental data, and does not try to define the accuracy of NMR ensembles for which methods such as ANSURR exist already [25].

The ability of AlphaFold2 to detect the presence of multiple conformations at the **residue level** does not necessarily extend to the full protein level, as it is not capable, when used in default mode, to detect proteins that can switch fold [63]. Likely this is because the overall fold prediction for the full protein is highly dependent on evolutionary information from a multiple sequence alignment, whereas at the residue level the overall atomic interaction information from the PDB that AlphaFold2 has learned is more likely to dominate. This is also reflected by the ShiftCrypt values in Fig. 4 F, with residues in the core and surrounding sheet regions, dependent on contacts between residues far removed from each other in the sequence, featuring higher pLDDT than core and surrounding helix, which are reliant on contacts between residues close to each other in the sequence. Notably, surrounding sheet, a relaxed conformational state with dynamic character, still has higher pLDDT than core helix, a rigid well-defined conformational state, showing that the prediction confidence of the former likely driven by evolutionary information for the overall fold of the protein, whereas the prediction for the latter is driven by highly local residue interactions.

In relation to actual experimentally observed dynamics, both estimated from NMR chemical shift data using the 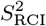 (Fig. 3 A/B) and ShiftCrypt methods (Fig. 4) and directly measured as *S*^2^ order parameters (Fig. 3 C/D), the pLDDT values show a binary relationship. Whereas high pLDDT values correspond to regions that have a stable, single rigid conformation in solution, residues with mid and lower pLDDT values do indicate the presence of multiple conformations and dynamics, but in this range the pLDDT value is only very weakly correlated with the degree of dynamics present. Whilst AlphaFold2 is thus capable of detecting a hard order/disorder boundary, as previously reported [15], it does not capture how dynamic a residue might be. This is not surprising given its training data, which mainly constitutes folded proteins organised in crystals and measured at cryogenic temperatures, so not capturing actual in solution dynamics information [16]. Residues observed to be dynamic in solution but still predicted with high pLDDT, on the other hand, seem to indicate regions that fold upon binding. This can be related to AlphaFold2 favoring the prediction of bound states of proteins, again due to its training data, where disordered regions would typically be missing in deposited PDB structures, and would only be visible when adopting a single conformation while interacting with another protein (or ligand [56]). The differences in secondary structure assignments between AlphaFold2 structure and NMR structures (Figs. 1, 2 and Supplementary Fig. 2) can also be partially explained by this observation.

The above discussion is summarised by the relation of pLDDT to the conformational state variability calculated from molecular dynamics trajectories (Fig. 6). High pLDDT values are spread out over low conformational state variability values, indicating little variation in conformational states, with said variation not picked up by AlphaFold2. Conversely, the mid to low pLDDT values have almost exclusively high conformational state variability, but again here the pLDDT does not pick up the degree of variation. These two trends highligh AlphaFold2’s capacity for binary order/disorder distinction, but its inability to capture the degree of conformational variability and dynamics.

Finally, this work shows that it is not trivial to estimate actual experimental dynamics from NMA on the AlphaFold2 models, with a complex relationship between these parameters. Whereas the rigid, well-folded parts of the protein are easily detected by the NMA RMSF, as they are by the pLDDT values, the interpretation of regions with lower pLDDT is dependent on whether these regions connect folded parts of the protein, with no straightforward relation between NMA RMSF and experimentally determined dynamics (Table 2, Fig. 8). This correlation improves when using NMR ensemble models as input for the NMA (Supplementary Figure 18). This contrasts with the ANSURR results, where rigidity is modelled as geometric constraints in protein structures using graph edges based on H-bonds and other interactions. Flexibility is computed by incrementally removing H-bond edges based on their energy thresholds (converted to Boltzmann population ratios) and observing when the C_α_ atom becomes flexible [25]. The increased accuracy with regard to this ANSURR rigidity score for AF2 models compared to NMR ensemble models is primarily attributed to AF2 models having more extensive and correctly placed H-bond networks, making them more rigid. The AlphaFold2 models then perform better in regions with extensive H-bond networks, with NMR structure ensembles rarely better except in dynamic regions [24]. The reliance of ANSURR on specific H-bonding networks, and not C_α_ atom distances as for NMA, might here be the key in explaining this difference. Whereas the variability between models in an NMR ensemble can interfere with the *in silico* definition of H-bonds at atomic precision, this variability does reflect in solution dynamics to some degree. NMA then seems to be better able to capture this information at the lower resolution of inter-C_α_ distances.

To conclude, the results in this work show that, whilst the AlphaFold2 pLDDT does indicate the presence or absence of multiple conformations and the associated protein dynamics, it does not capture the gradations of dynamics nor the number of possible conformations present. The RMSF of NMA on the AlphaFold2 models equally does not capture such information. Experimental data, such as from NMR, and more fine-grained computational approaches, such as molecular dynamics simulations, therefore remain invaluable to assess the movements and conformational states of proteins. While AlphaFold2 has fast-forwarded the field of structural biology and our understanding of the space of protein folds, its limitations are, as with any machine learning method, determined by its training data, which does not incorporate dynamics. Notwithstanding the applicability of AlphaFold2 on predicting multiple conformations on a selected set of well-studied proteins [7], this study highlights the complexity of the problem when relating AlphaFold2 to large scale experimental data capturing the presence of multiple conformations and the degree of dynamics present. Our ability to predict multiple conformations and dynamics will therefore likely remain limited until the lack of reliable and extensive experimental training data that encompasses multiple conformational states and the dynamics of proteins is resolved.

## Supporting information

Supplementary Information part 1

Supplementary Information part 2

## Acknowledgements

This work has been supported by the European Union’s Horizon 2020 research and innovation program under the Marie Skłodowska-Curie grant agreement [813239 to J.G.-G. and A.D.]; Research Foundation Flanders (FWO) International Research Infrastructure [I000323N to W.V. and G.0328.16N to A.D.]; COST Action ML4NGP, CA21160, supported by COST (European Cooperation in Science and Technology). We also acknowledge project funding to A.G. of the FWO (project G002520N and project G094023N), BOF of Ghent University, and the European Union (ERC, 101086145 PASTIME). The computational resources (Stevin Supercomputer Infrastructure) and services used in this work were provided by the VSC (Flemish Supercomputer Center), funded by Ghent University, FWO and the Flemish Government – department EWI.

## Author Contributions

Conceptualization: WV. Methodology: JGG, BD, WV, AG. Software: JGG, BD. Formal analysis: JGG, BD. Web server and interactive analysis: JGG, AD. Data Curation: JGG, BD, WV, AG. Writing: JGG, BD, WV, AG. Visualisation: JGG, BD. Supervision: WV, AG. Funding acquisition: WV, AG.

