## Supplementary Information part 1 for "Gradations in protein dynamics captured by experimental NMR are not well represented by AlphaFold2 models and other computational metrics"

July 17, 2024

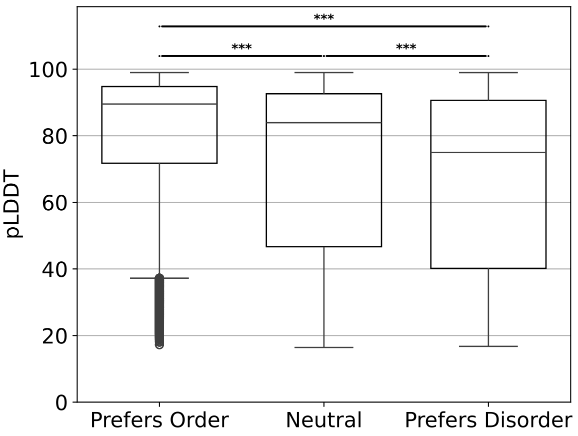


Supplementary Figure 1: **pLDDT values per amino acid order classes.** The amino acids from the $S^2_{\mathrm{RCI}}$ dataset were clustered in 3 classes according to their order preference: preferentially ordered (cysteine, phenylalanine, isoleucine, leucine, valine, tryptophan & tyrosine; N=106,363), neutral (alanine, glutamic acid, lysine, methionine, glutamine, arginine, threonine & histidine; N=155,050) and preferentially disordered (aspartic acid, glycine, asparagine, proline & serine; N=112,945). The distributions were tested with a Mann–Whitney U test, which resulted in p-values $< 0.001$ in all tests.

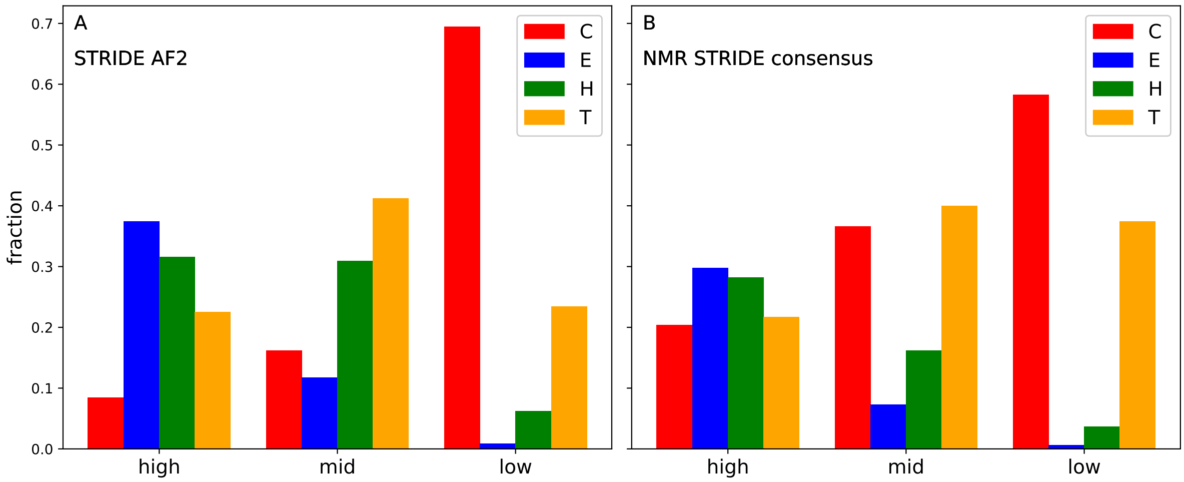


Supplementary Figure 2: **Abundance of secondary structures by pLDDT ranges.** A) The STRIDE secondary structure assignment from the structure that AlphaFold2 produces. B) The STRIDE consensus secondary structure assignment here provided is the most abundantly assigned secondary structure in the ensemble of NMR structures available. The STRIDE consensus assignment (panel B) produces a decrease in abundance for helix (H) and sheet (E) fractions as the pLDDT decreases, as well as an increase in coil (C) and turn (T) conformations. These tendencies are maintained for AlphaFold2's single structure assignment (panel A) in sheet and coil fractions, but not for helix and turn.

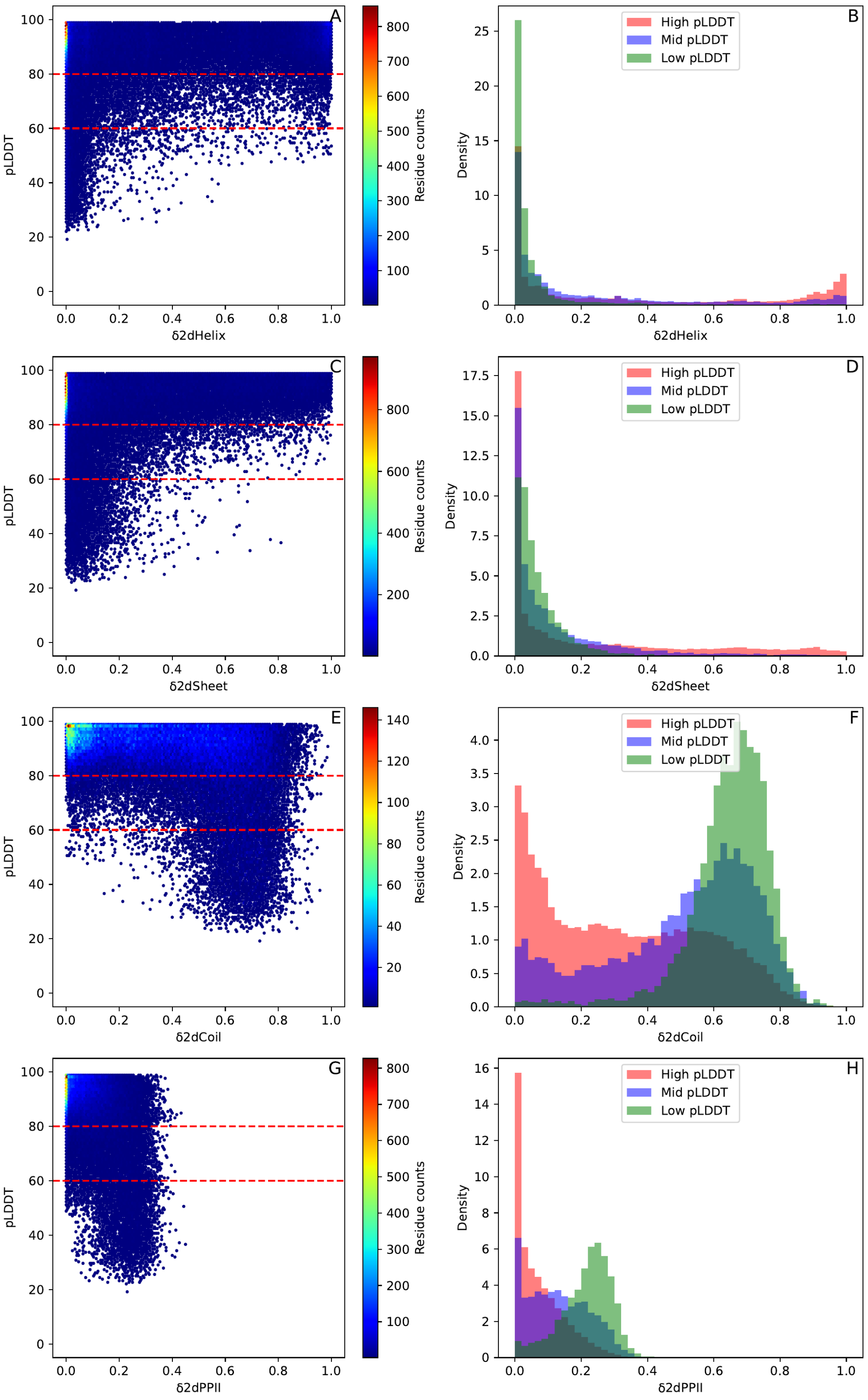


Supplementary Figure 3: **Comparison of pLDDT and $\delta 2D$ populations.** For all available residues in the $S^2_{\mathrm{RCI}}$ dataset, the populations of conformational states were calculated with $\delta$2D method. High populations values of a conformation indicate high presence of such conformation in the ensemble, and vice-versa. A-D: For both Helix and Sheet conformations, higher $\delta$2D populations are obtained for residues featuring high pLDDT ($\geq 80$, N=62,014), gradually adopting lower populations for mid ($80 >$ pLDDT $\geq 60$, N=8,539) and low ($< 60$, N=4,773) pLDDT values. E-H: Coil and PPII populations are prominently higher for higher for low pLDDT ranges than for mid and high ranges. Mann-Whitney two-sided U tests p-values between each pLDDT-stratified distribution for each $\delta 2D$ conformation confirmed difference at p-values $< 0.001$ (table 2).

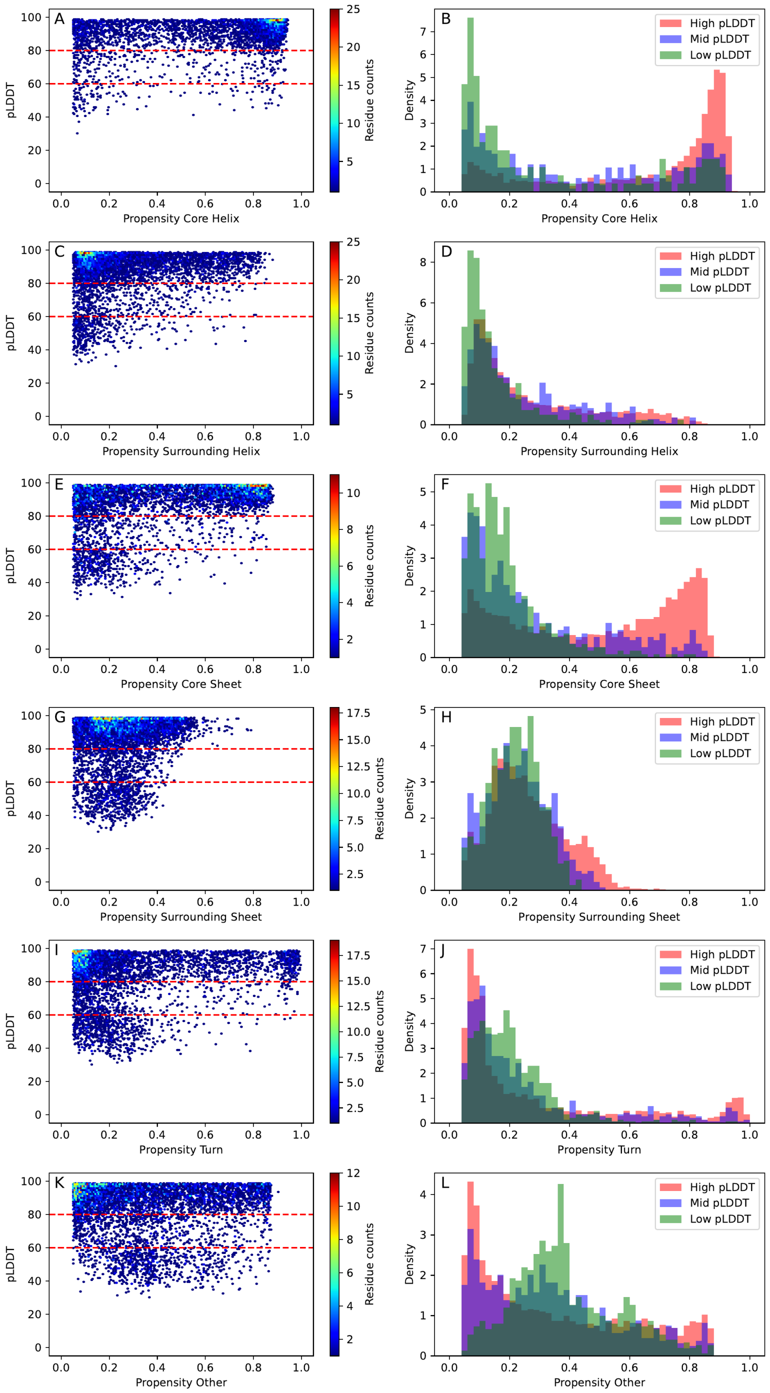


Supplementary Figure 4: **Comparison between AlphaFold2 pLDDT and Constava's conformational state propensities.** The pLDDT of each residue was plotted against the propensity for each of the 6 conformational states calculated in Constava. Any propensity below 0.05 was deemed non-informative and discarded to facilitate interpretation. All residues in the MD dataset were stratified in high (N=9,523), mid (N=1,038) and low (N=809) AlphaFold2 ranges. A & B) Hexagonal binning of AlphaFold3 C-$\alpha$ pLDDT vs Core Helix propensity and its associated pLDDT-stratified distributions. C & D) Hexagonal binning of AlphaFold3 C-$\alpha$ pLDDT vs Surrounding Helix propensity and its associated pLDDT-stratified distributions. E & F) Hexagonal binning of AlphaFold3 C-$\alpha$ pLDDT vs Core Sheet propensity and its associated pLDDT-stratified distributions. G & H) Hexagonal binning of AlphaFold3 C-$\alpha$ pLDDT vs Surrounding Sheet propensity and its associated pLDDT-stratified distributions. I & J) Hexagonal binning of AlphaFold3 C-$\alpha$ pLDDT vs Turn propensity and its associated pLDDT-stratified distributions. K & L) Hexagonal binning of AlphaFold3 C-$\alpha$ pLDDT vs Other propensity and its associated pLDDT-stratified distributions. The p-values for the Mann-Whitney two-sided U tests between each pLDDT-stratified distribution for every conformational state can be found in supplementary table 1.

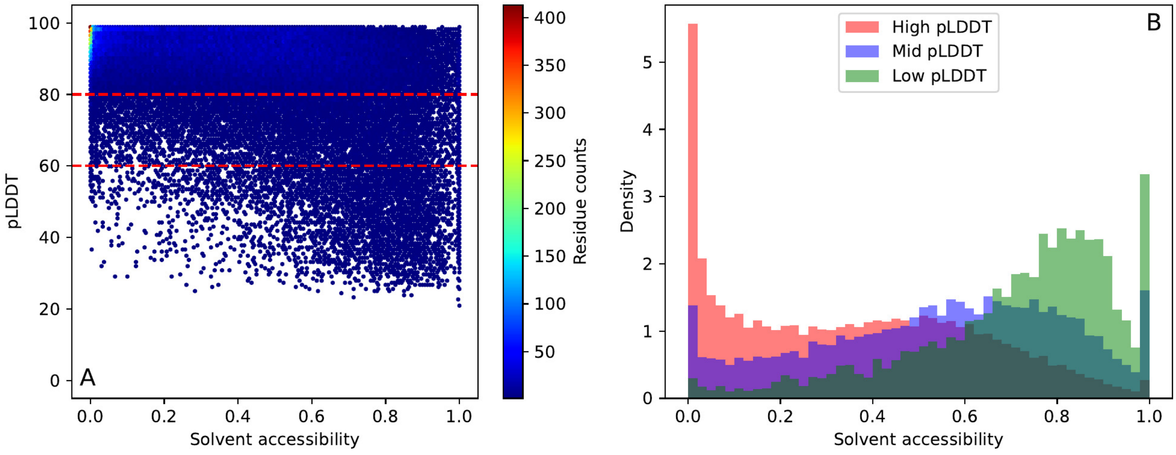


Supplementary Figure 5: **Comparison of pLDDT and solvent accessibility.** A) The AlphaFold2 pLDDT in the $S^2_{RCI}$ dataset were plotted against the solvent accessibility score for every residue with available values, in an hexagonal binning plot. B) Distributions of solvent accessibility scores, stratified in high ($\geq$ 80, N=62,319), mid (80 $>$ pLDDT $\geq$ 60, N=8,708) and low ($<$ 60, N=4,842) AlphaFold2 pLDDT values. Mann-Whitney two-sided U test confirmed significant differences between all distribution pairs with a p-value $<$ 0.001 (table 3).

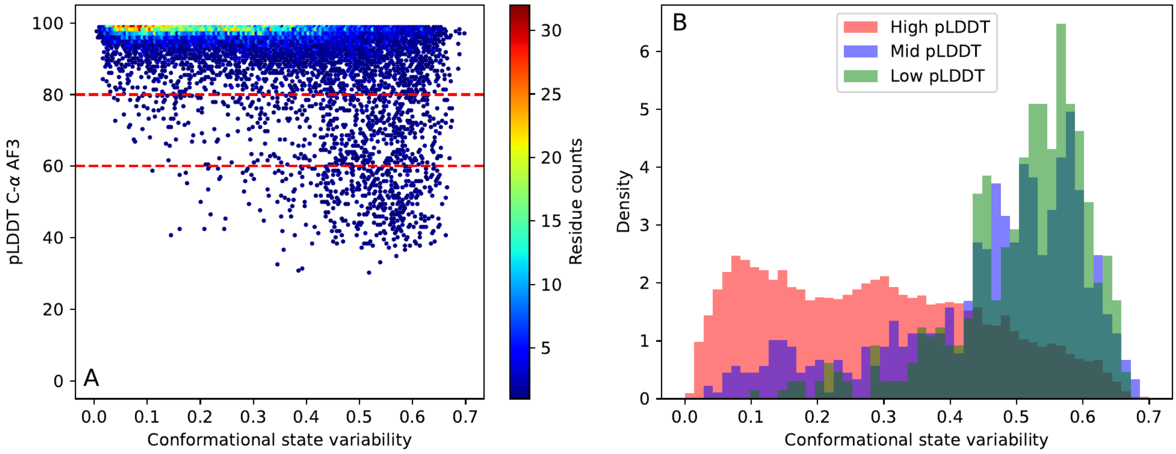


Supplementary Figure 6: **AlphaFold3 C-$\alpha$ pLDDT vs. conformational state variability.** A) High pLDDT values ($\geq$ 80, N = 10,272) concentrate in areas with lower conformational state variability. Low pLDDT $<$ 60, N = 463) usually correspond to residues with high conformational state variability. B) This tendency is more clearly observed with pLDDT-stratified distributions, which shows that low pLDDT residues correspond to residues with high conformational state variability, therefore with high potential to exist in multiple conformations, and vice-versa for high pLDDT and low variability residues. Mid pLDDT residues 80 $>$ pLDDT $\geq$ 60, N = 634) exhibit an intermediate distribution. Mann-Whitney two-sided U test yielded a p-value $<$ 0.001 between all distributions (table 3). The associated distributions per propensity can be found in supplementary fig. 7.

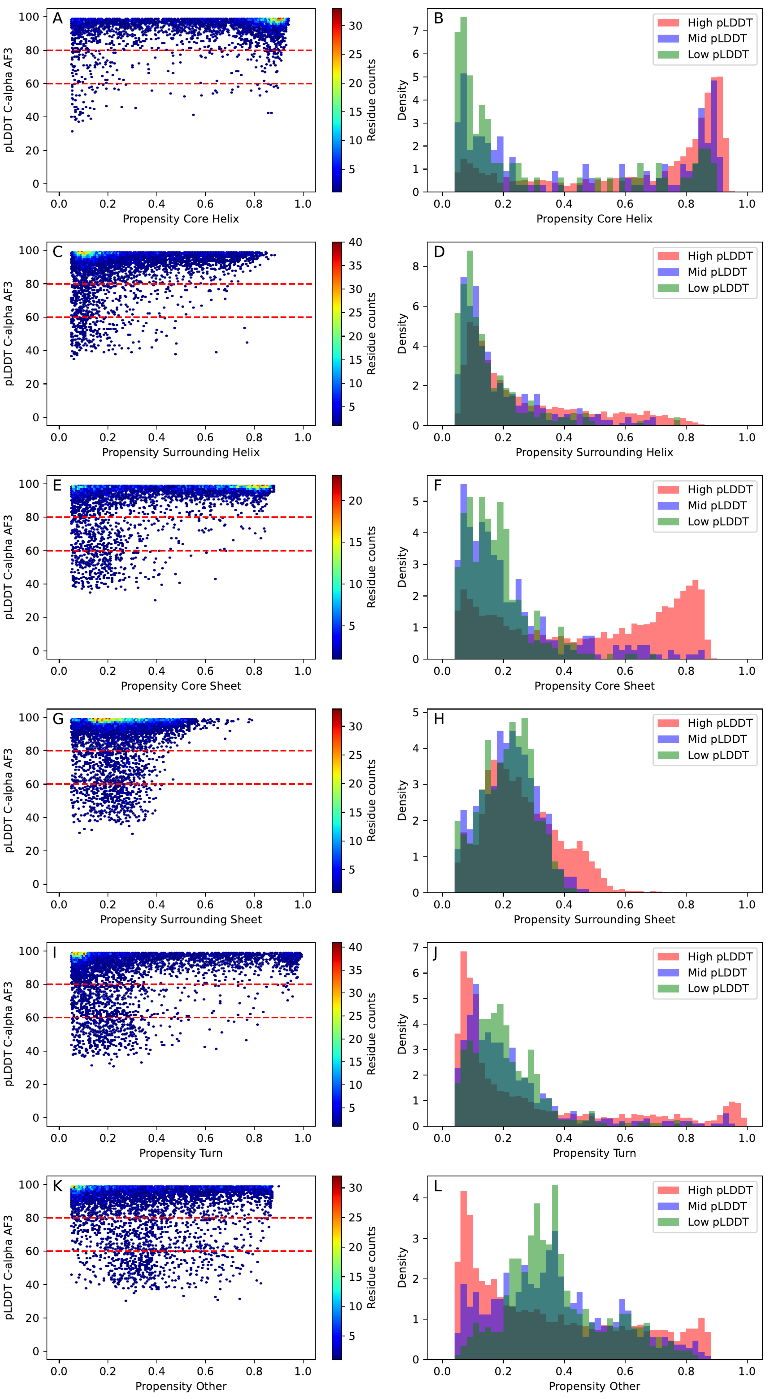


Supplementary Figure 7: **Comparison between AlphaFold3 C-$\alpha$ pLDDT and Constava's conformational state propensities.** The pLDDT of each residue was plotted against the propensity for each of the 6 conformational states calculated in Constava. Any propensity below 0.05 was deemed non-informative and discarded to facilitate interpretation. All residues in the MD dataset were stratified in high (N=10,272), mid (N=634) and low (N=463) AlphaFold3 C-$\alpha$ ranges. A & B) Hexagonal binning of AlphaFold3 C-$\alpha$ pLDDT vs Core Helix propensity and its associated pLDDT-stratified distributions. C & D) Hexagonal binning of AlphaFold3 C-$\alpha$ pLDDT vs Surrounding Helix propensity and its associated pLDDT-stratified distributions. E & F) Hexagonal binning of AlphaFold3 C-$\alpha$ pLDDT vs Core Sheet propensity and its associated pLDDT-stratified distributions. G & H) Hexagonal binning of AlphaFold3 C-$\alpha$ pLDDT vs Surrounding Sheet propensity and its associated pLDDT-stratified distributions. I & J) Hexagonal binning of AlphaFold3 C-$\alpha$ pLDDT vs Turn propensity and its associated pLDDT-stratified distributions. K & L) Hexagonal binning of AlphaFold3 C-$\alpha$ pLDDT vs Other propensity and its associated pLDDT-stratified distributions. The p-values for the Mann-Whitney two-sided U tests between each pLDDT-stratified distribution for every conformational state can be found in supplementary table 1

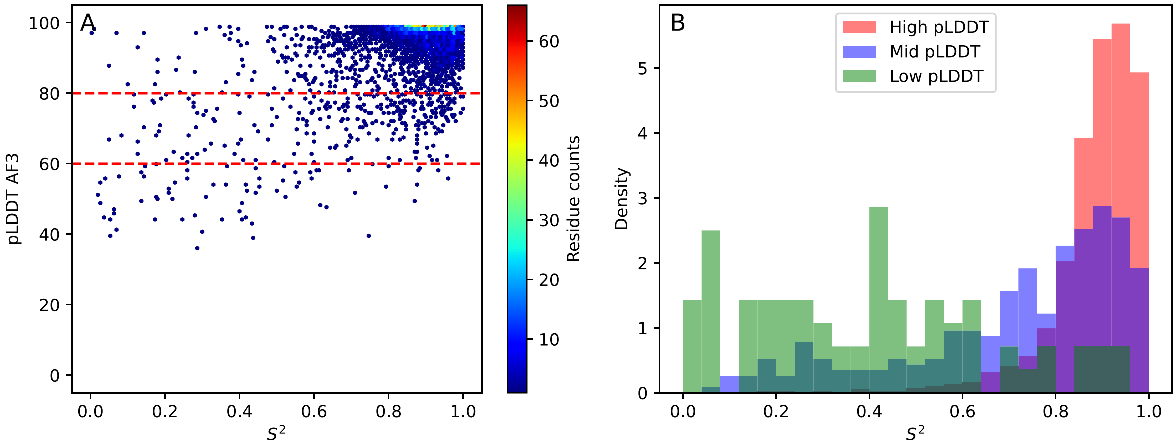


Supplementary Figure 8: **Comparison between AlphaFold3 C-$\alpha$ pLDDT and $S^2$.** Most residues in this dataset exhibit high pLDDT and high $S^2$ values, represented by warmer hues in panel A. Due to the limited and uneven residues sample sizes (high pLDDT, $\geq 80$, N = 4,125; mid pLDDT, $80 > \text{pLDDT} \geq 60$, N = 287; low pLDDT, $< 60$, N = 70), it is challenging to make definitive conclusions about the sparsely populated mid and low pLDDT regions. Panel B illustrates the distributions of $S^2$ values of each pLDDT range, for which Mann-Whitney two-sided U test yielded a p-value $< 0.001$ between all distributions (table 3).

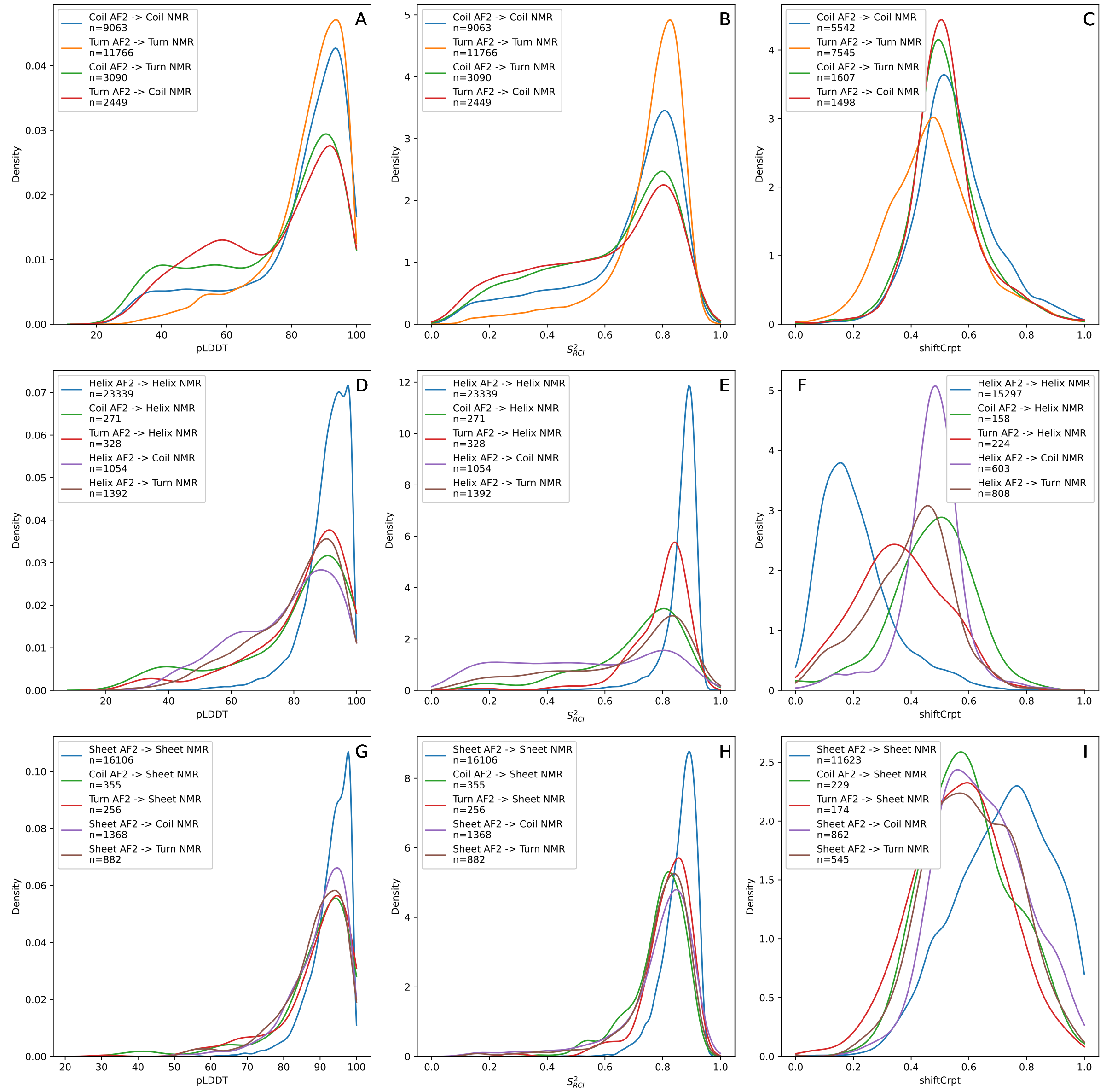


Supplementary Figure 9: **Distribution of STRIDE fold assignment matches and mismatches between AlphaFold2 structures and consensus NMR ensemble assignments, for diverse metrics.** Those residues in the $S^2_{RCI}$ dataset with STRIDE consensus were stratified according to their secondary assignment pairs, derived from AlphaFold2 structures and the NMR models consensus assignment. A, D & G) pLDDT distributions for residues whose consensus STRIDE assignment from NMR ensembles and from AlphaFold2 models match or mismatch. B, E & H) $S^2_{RCI}$ distributions for residues whose consensus STRIDE assignment from NMR ensembles and from AlphaFold2 models match or mismatch. C, F & I) shiftCrypt distributions for residues whose consensus STRIDE assignment from NMR ensembles and from AlphaFold2 models match or mismatch.

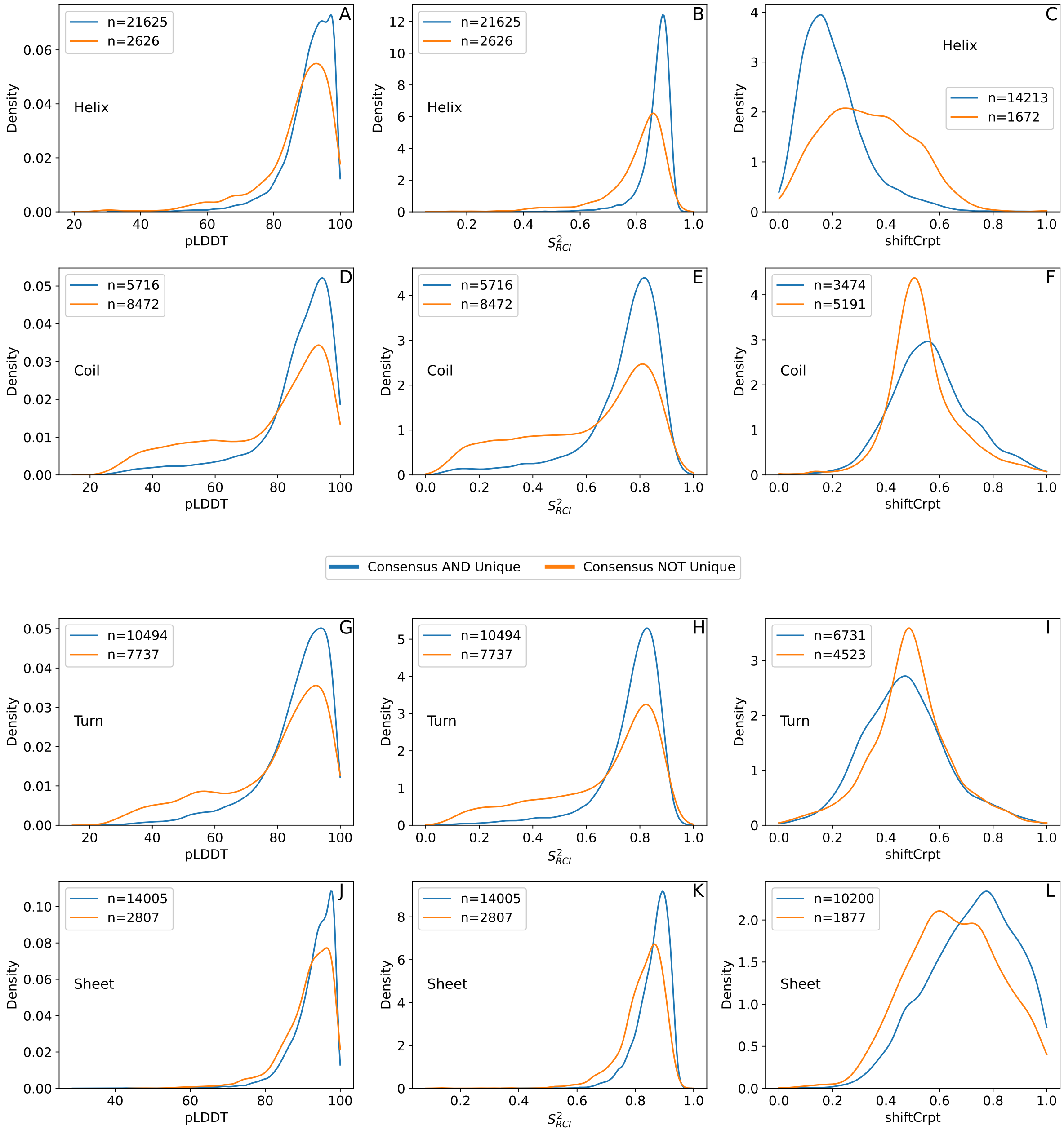


Supplementary Figure 10: **Distribution of residues with NMR consensus STRIDE assignment with and without unique STRIDE assignment, for diverse metrics.** Those residues in the $S^2_{RCI}$ dataset with STRIDE consensus were stratified according to whether they featured a unique STRIDE assignment across all the models in their corresponding NMR ensemble. A, D, G & J) pLDDT distributions for $\alpha$-helix, coil, turn and $\beta$-sheet respectively. B, E, H & K) $S^2_{RCI}$ distributions for $\alpha$-helix, coil, turn and $\beta$-sheet respectively. C, F, I & L) shiftCrypt distributions for $\alpha$-helix, coil, turn and $\beta$-sheet respectively.

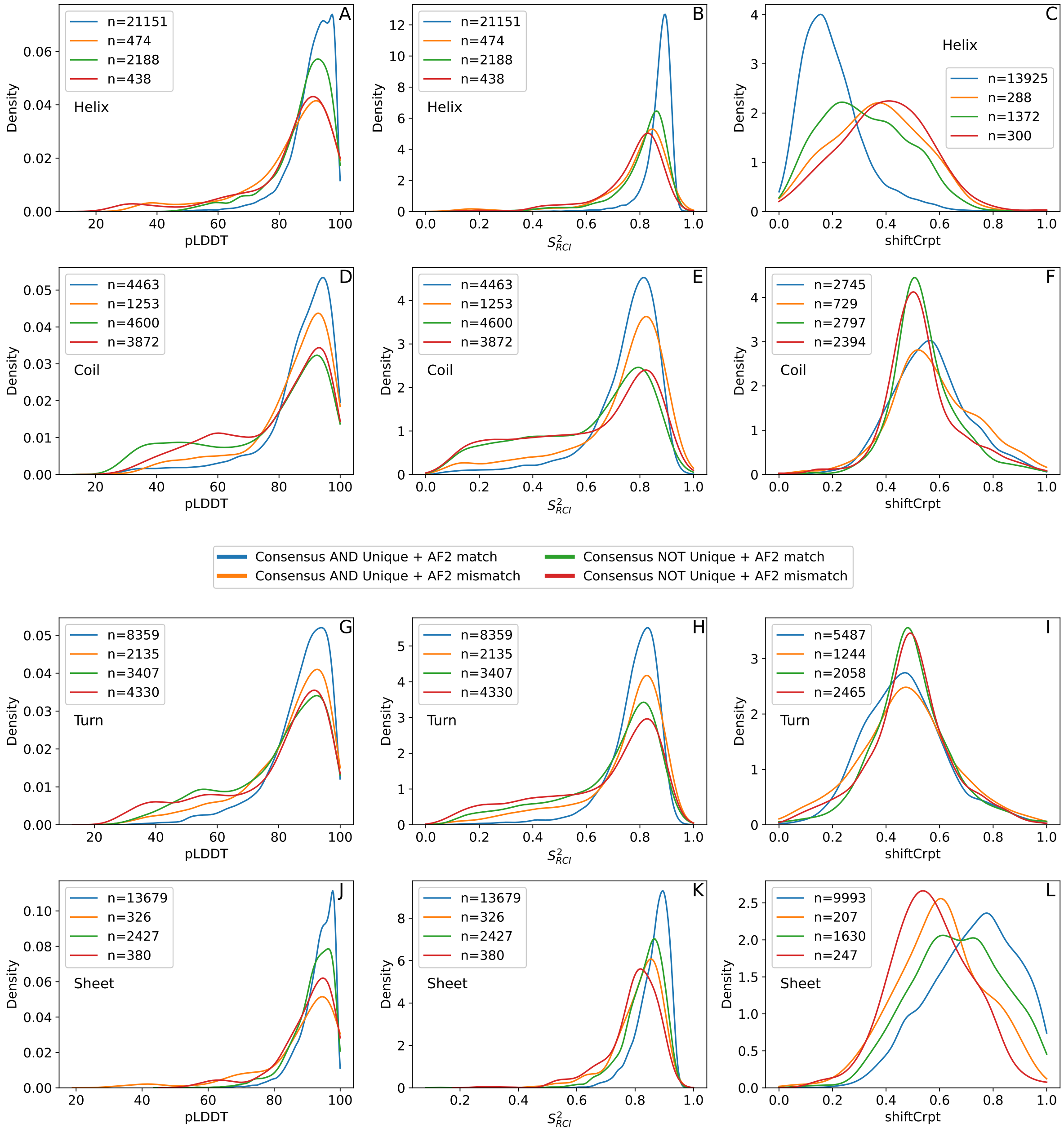


Supplementary Figure 11: **Distribution of residues with NMR consensus STRIDE assignment with and without unique STRIDE assignment, with matching and mismatching NMR and AlphaFold2 fold assignments, for diverse metrics.** Those residues in the $S^2_{RCI}$ dataset with STRIDE consensus were stratified according to whether they featured a unique STRIDE assignment across all the models in their corresponding NMR ensemble. Then, they were further stratified on whether or not their NMR consensus STRIDE assignment matched the AlphaFold2 STRIDE assignment. A, D, G & J) pLDDT distributions for $\alpha$-helix, coil, turn and $\beta$-sheet respectively. B, E, H & K) $S^2_{RCI}$ distributions for $\alpha$-helix, coil, turn and $\beta$-sheet respectively. C, F, I & L) shiftCrypt distributions for $\alpha$-helix, coil, turn and $\beta$-sheet respectively.

Supplementary Table 1: **Results of the Mann-Whitney two-sided U tests between pLDDT-stratified subsets per Constava's conformational states on the MD dataset.** For each of Constava's conformational states, the pLDDT-stratified subsets were tested against each other with a Mann-Whitney two-sided U test to assess differential distributions.

| Conformational state | AlphaFold version | pLDDT ranges | Difference between means | p-value |
|---|---|---|---|---|
| Core Helix | 2 | High-Mid | 0.2161 | $6.67 \times 10^{-34}$ |
| Core Helix | 2 | High-Low | 0.3427 | $2.52 \times 10^{-31}$ |
| Core Helix | 2 | Mid-Low | 0.1266 | $1.53 \times 10^{-5}$ |
| Surr. Helix | 2 | High-Mid | 0.0264 | 0.01955 |
| Surr. Helix | 2 | High-Low | 0.1168 | $5.17 \times 10^{-43}$ |
| Surr. Helix | 2 | Mid-Low | 0.0903 | $1.24 \times 10^{-20}$ |
| Core Sheet | 2 | High-Mid | 0.2254 | $1.60 \times 10^{-60}$ |
| Core Sheet | 2 | High-Low | 0.3074 | $2.12 \times 10^{-104}$ |
| Core Sheet | 2 | Mid-Low | 0.0821 | $1.58 \times 10^{-5}$ |
| Surr. Sheet | 2 | High-Mid | 0.0339 | $1.96 \times 10^{-8}$ |
| Surr. Sheet | 2 | High-Low | 0.0455 | $3.02 \times 10^{-15}$ |
| Surr. Sheet | 2 | Mid-Low | 0.0117 | 0.08711 |
| Turn | 2 | High-Mid | 0.0394 | 0.44796 |
| Turn | 2 | High-Low | 0.0826 | 0.22141 |
| Turn | 2 | Mid-Low | 0.0432 | 0.45743 |
| Other | 2 | High-Mid | 0.0082 | 0.28323 |
| Other | 2 | High-Low | -0.0397 | $6.19 \times 10^{-15}$ |
| Other | 2 | Mid-Low | -0.0479 | $1.85 \times 10^{-9}$ |
| Core Helix | 3 | High-Mid | 0.1990 | $4.07 \times 10^{-14}$ |
| Core Helix | 3 | High-Low | 0.3498 | $4.63 \times 10^{-20}$ |
| Core Helix | 3 | Mid-Low | 0.1507 | $1.85 \times 10^{-4}$ |
| Surr. Helix | 3 | High-Mid | 0.0861 | $6.18 \times 10^{-18}$ |
| Surr. Helix | 3 | High-Low | 0.1203 | $7.55 \times 10^{-27}$ |
| Surr. Helix | 3 | Mid-Low | 0.0343 | 0.00256 |
| Core Sheet | 3 | High-Mid | 0.2657 | $1.94 \times 10^{-58}$ |
| Core Sheet | 3 | High-Low | 0.3019 | $6.68 \times 10^{-64}$ |
| Core Sheet | 3 | Mid-Low | 0.0362 | 0.29869 |
| Surr. Sheet | 3 | High-Mid | 0.0423 | $6.90 \times 10^{-10}$ |
| Surr. Sheet | 3 | High-Low | 0.0462 | $1.51 \times 10^{-10}$ |
| Surr. Sheet | 3 | Mid-Low | 0.0039 | 0.66095 |
| Turn | 3 | High-Mid | 0.0628 | 0.48281 |
| Turn | 3 | High-Low | 0.0779 | 0.13434 |
| Turn | 3 | Mid-Low | 0.0152 | 0.33197 |
| Other | 3 | High-Mid | -0.0178 | 0.000078 |
| Other | 3 | High-Low | -0.0288 | $2.72 \times 10^{-8}$ |
| Other | 3 | Mid-Low | -0.0110 | 0.15908 |

Supplementary Table 2: **Results of the Mann-Whitney two-sided U tests between pLDDT-stratified subsets for $S^2_{RCI}$ $\delta 2D$ conformations in AlphaFold2.** For each conformation type, the pLDDT-stratified subsets were tested against each other with a Mann-Whitney two-sided U test to assess differential distributions. *Note: p-values marked with * were too low for Scipy to differentiate from 0.*

| Conformation | pLDDT ranges | Difference between means | p-value |
|---|---|---|---|
| $\delta 2D$ Helix | High-Mid | 0.1185 | $3.38 \times 10^{-55}$ |
| $\delta 2D$ Helix | High-Low | 0.2896 | 0.0* |
| $\delta 2D$ Helix | Mid-Low | 0.1711 | $1.23 \times 10^{-300}$ |
| $\delta 2D$ Sheet | High-Mid | 0.1181 | $5.03 \times 10^{-49}$ |
| $\delta 2D$ Sheet | High-Low | 0.1723 | $5.44 \times 10^{-53}$ |
| $\delta 2D$ Sheet | Mid-Low | 0.0542 | $2.63 \times 10^{-7}$ |
| $\delta 2D$ Coil | High-Mid | -0.1775 | 0.0* |
| $\delta 2D$ Coil | High-Low | -0.3184 | 0.0* |
| $\delta 2D$ Coil | Mid-Low | -0.1410 | $3.76 \times 10^{-305}$ |
| $\delta 2D$ PPII | High-Mid | -0.0592 | 0.0* |
| $\delta 2D$ PPII | High-Low | -0.1435 | 0.0* |
| $\delta 2D$ PPII | Mid-Low | -0.0843 | 0.0* |

Supplementary Table 3: **Results of the Mann-Whitney two-sided U tests between pLDDT-stratified subsets.** For each dataset, the pLDDT-stratified subsets were tested against each other with a Mann-Whitney two-sided U test to assess differential distributions. *Note: p-values marked with * were too low for Scipy to differentiate from 0.*

| Dataset | Metric | AlphaFold version | pLDDT ranges | Difference between means | p-value |
|---|---|---|---|---|---|
| $S^2_{\mathrm{RCI}}$ | $S^2_{\mathrm{RCI}}$ | 2 | high-mid | 0.1417 | 0* |
| $S^2_{\mathrm{RCI}}$ | $S^2_{\mathrm{RCI}}$ | 2 | high-low | 0.4150 | 0* |
| $S^2_{\mathrm{RCI}}$ | $S^2_{\mathrm{RCI}}$ | 2 | mid-low | 0.2732 | 0* |
| $S^2_{\mathrm{RCI}}$ | ShiftCrypt | 2 | high-mid | 0.0152 | 0.002 |
| $S^2_{\mathrm{RCI}}$ | ShiftCrypt | 2 | high-low | -0.0201 | $4.55 \times 10^{-7}$ |
| $S^2_{\mathrm{RCI}}$ | ShiftCrypt | 2 | mid-low | -0.0353 | $9.45 \times 10^{-21}$ |
| $S^2_{\mathrm{RCI}}$ | Solvent accessibility | 2 | high-mid | -0.1851 | 0* |
| $S^2_{\mathrm{RCI}}$ | Solvent accessibility | 2 | high-low | -0.3521 | 0* |
| $S^2_{\mathrm{RCI}}$ | Solvent accessibility | 2 | mid-low | -0.1670 | $1.93 \times 10^{-297}$ |
| $S^2$ | $S^2$ | 2 | high-mid | 0.1352 | $5.60 \times 10^{-24}$ |
| $S^2$ | $S^2$ | 2 | high-low | 0.4238 | $1.19 \times 10^{-53}$ |
| $S^2$ | $S^2$ | 2 | mid-low | 0.2886 | $2.12 \times 10^{-19}$ |
| $S^2$ | $S^2$ | 3 | high-mid | 0.1681 | $1.74 \times 10^{-43}$ |
| $S^2$ | $S^2$ | 3 | high-low | 0.4904 | $6.36 \times 10^{-38}$ |
| $S^2$ | $S^2$ | 3 | mid-low | 0.3223 | $2.45 \times 10^{-16}$ |
| MD | Conf. state var. | 2 | high-mid | -0.1468 | $3.46 \times 10^{-143}$ |
| MD | Conf. state var. | 2 | high-low | -0.2197 | $1.23 \times 10^{-246}$ |
| MD | Conf. state var. | 2 | mid-low | -0.0729 | $1.48 \times 10^{-22}$ |
| MD | Conf. state var. | 3 | high-mid | -0.1743 | $1.46 \times 10^{-126}$ |
| MD | Conf. state var. | 3 | high-low | -0.2258 | $4.6 \times 10^{-155}$ |
| MD | Conf. state var. | 3 | mid-low | -0.0515 | $2.08 \times 10^{-7}$ |
