## Supplementary Information part 2 for "Gradations in protein dynamics captured by experimental NMR are not well represented by AlphaFold2 models and other computational metrics"

On the $S^2_{\mathrm{RCI}}$ dataset of 762 proteins, WEBnma was carried out on the 762 AlphaFold2 models. Therefore, based on WEBnma output, the final dataset consists of 762 AlphaFold2 models. The predicted RMSF values of each coil residue in all 762 proteins are shown in (Supplemental Figure 12). The Supplemental Figure shows some extreme RMSF values in low-pLDDT regions. As explained in the main text, these extreme values can originate artificially from loosely packed stretches in the protein structure. We have therefore adapted the RMSF analysis to reduce the artificial RMSF outliers. As shown in the following (section Truncation criterion), the N- and C-terminal tails from AlphaFold2 models were truncated. The truncation criterion is based on the number of C$\alpha$ contacts. The final number of truncated proteins in the dataset was 755, and the remaining 7 did not require cutting of termini. Subsequently, normal mode analysis with WEBnma was again performed on these 755 truncated models, and the RMSF was recomputed. The RMSF results are shown for 762 proteins, including both the 755 truncated models and the 7 models that did not require termini cutting (referred to as truncated $S^2_{\mathrm{RCI}}$ dataset). Apart from (Supplemental Figure 12, Supplemental Figure 13, and Supplemental Table 4) all figures and data in the main document and SI contain the RMSF of truncated $S^2_{\mathrm{RCI}}$ dataset. Supplemental Figure 13 shows the effect of the truncation by comparing the RMSF before truncation and the RMSF after truncation.

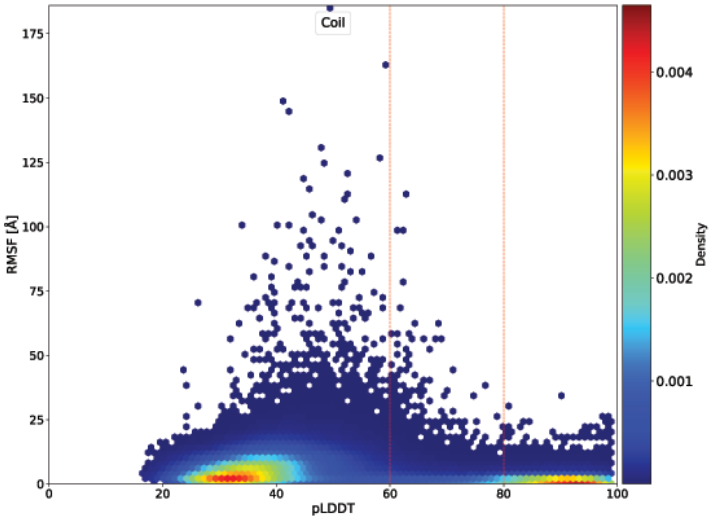


Supplemental Figure 12 **Comparison of pLDDT and RMSF in coils** pLDDT vs RMSF of 762 proteins for coil residues before truncation. The colour bar represents the Gaussian kernel density estimate of the dataset. The red vertical lines divide the dataset into high pLDDT ($\geq 80$), mid ($\geq 60$ pLDDT $< 80$) and low ($< 60$) pLDDT regions.

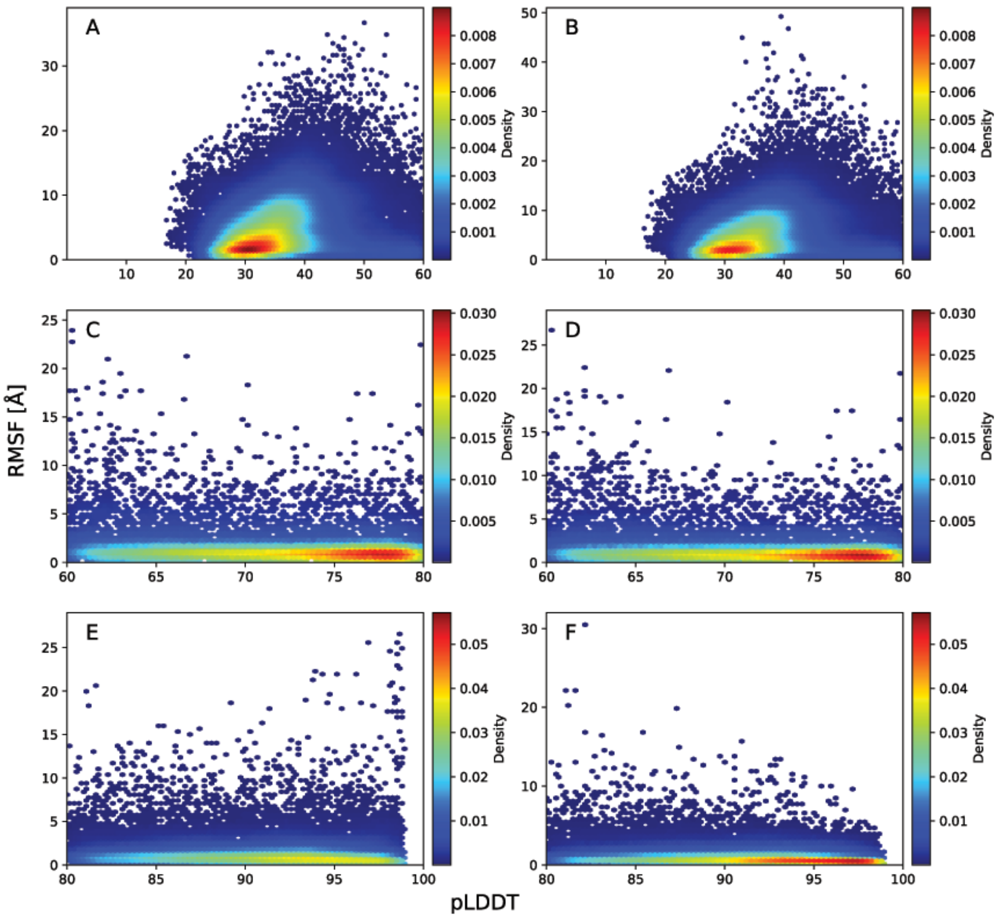


Supplemental Figure 13 **Comparison of pLDDT and RMSF in coils** RMSF vs pLDDT of amino acid residues exhibiting coils in non-truncated (A, C, E) and truncated (B, D, F) AlphaFold2 structures in low-pLDDT (A, B), mid-pLDDT (C, D), and c) high-pLDDT (E, F) regions. Only the amino acids that are present in both the non-truncated and truncated AlphaFold2 models are included.

Supplemental Table 4 RMSF values of are grouped according to pLDDT in low-pLDDT, mid-pLDDT and high-pLDDT for AlphaFold2 models before truncation. The table reports the minimum, maximum, mean, and standard deviation for each group.

| | low-pLDDT | | | | mid-pLDDT | | | | high-pLDDT | | | |
|---|---|---|---|---|---|---|---|---|---|---|---|---|
| **secondary structure** | **min** | **max** | **mean** | **std** | **min** | **max** | **mean** | **std** | **min** | **max** | **mean** | **std** |
| Coil | 0.22 | 184.99 | 6.28 | 5.83 | 0.21 | 112.52 | 3.00 | 5.16 | 0.15 | 33.97 | 1.58 | 1.84 |
| Strand | 0.28 | 8.05 | 1.81 | 1.66 | 0.23 | 21.63 | 1.49 | 1.6 | 0.15 | 29.67 | 1.35 | 1.49 |
| α-Helix | 0.24 | 25.98 | 2.11 | 2.15 | 0.18 | 23.61 | 2.02 | 2.37 | 0.14 | 24.68 | 1.57 | 1.87 |
| Turn | 0.22 | 37.15 | 2.62 | 2.74 | 0.20 | 33.05 | 2.04 | 2.56 | 0.15 | 32.52 | 1.63 | 1.91 |
| $3_{10}$-Helix | 0.29 | 27.83 | 2.27 | 3.29 | 0.21 | 27.39 | 2.11 | 3.07 | 0.20 | 30.04 | 1.37 | 1.55 |
| Bridge | 0.31 | 19.52 | 1.78 | 2.50 | 0.26 | 13.06 | 1.66 | 2.04 | 0.15 | 29.29 | 1.48 | 1.78 |

### Truncation criterion

For determining N- and/or C-termini truncation, the C$\alpha$ contacts were assessed within a 10 Å (1 nm) cut-off for each protein in the dataset. In proteins, helices and strand consistently exhibited significant contacts, surpassing approximately 13 contacts per residue across the dataset with lower RMSF (<20 Å) as shown in (Supplemental Figure 14) An example is shown in Supplemental Figure 15. In contrast, coils showed fewer than 13 contacts per residue and showed very high RMSF (>50 Å). Thus, a 13-contact cutoff was selected to truncate the termini.

Following this criterion, all first residues with fewer than 13 contacts were cut both in the N-terminal and C-terminal. Consequently, if the first residue of an N- or C-terminal has ≥ 13 contacts, this terminal was not truncated. Only the termini were truncated, so an accidental low contact region in the core of the protein would not get cut.

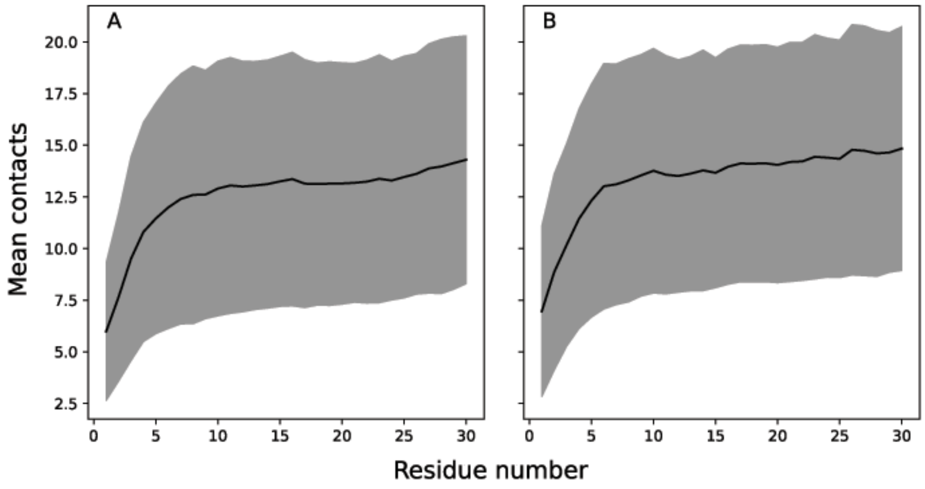


Supplemental Figure 14 **Mean number of contacts** Mean number of C$\alpha$ contacts within a 10 Åcut-off for first 30 residues depicting N-terminal (A) and last 30 residues depicting C-terminal (B) averaged over all 762 proteins. The black line represents the mean contacts, with the standard deviation shown as grey shaded area.

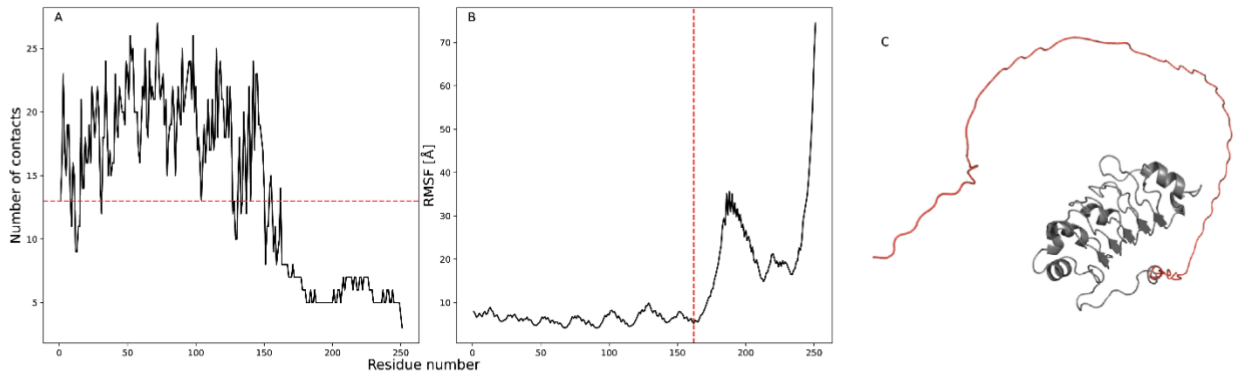


Supplemental Figure 15 **Number of C$\alpha$ contacts profile** (A) and RMSF profile (B) of Q92688. The red dashed line in A represents the contact cut-off (13 contacts) and red dashed line in B represents the RMSF at contact cut-off. The 3D structure of Q92688 is shown in C with a red highlighted region for truncation.

### Additional analysis of RMSF and correlation with pLDDT or $S^2_{RCI}$

The RMSF values for the dataset with truncated dataset are further analysed (now 762 proteins) according to their secondary structure element as predicted by STRIDE.

The six considered secondary structure elements are coil, strand, $\alpha$-helix, turn, $3_{10}$-helix, and bridge. The tables report the minimum, maximum, mean, and standard deviation for the RMSF values in each secondary structure group. Supplemental Table 5 gives three columns according to the pLDDT value as given by AlphaFold2: low-pLDDT, mid-pLDDT, and high-pLDDT. Supplemental Table 6 gives three columns according to the $S^2_{\mathrm{RCI}}$ value as included in the truncated $S^2_{\mathrm{RCI}}$ dataset: flexible, ambiguous, and rigid.

Next, the Pearson correlation coefficient between the RMSF values and the pLDDT values were computed in each group (low-pLDDT, mid-pLDDT, and high-pLDDT) in (Supplemental Table 7). Moreover, the Pearson correlation coefficient between RMSF and pLDDT was computed, without considering the subgroups of pLDDT (Supplemental Table 7). Similarly, the Pearson correlation coefficient between RMSF and $S^2_{\mathrm{RCI}}$ was computed for each group (flexible, ambiguous, and rigid), and without considering subgroups of $S^2_{\mathrm{RCI}}$ (Supplemental Table 7). For both RMSF and pLDDT, RMSF and $S^2_{\mathrm{RCI}}$, the Pearson correlation was calculated for each secondary structure group, and without the classification of secondary structure.

Next, The Pearson correlation coefficient between the RMSF values and $S^2_{\mathrm{RCI}}$ can also be computed for each individual AlphaFold2 and NMR model in the truncated $S^2_{\mathrm{RCI}}$ dataset. This is reported as a histogram in Supplemental Figure 18 (blue) using the RMSF values of the 746 AlphaFold2 models and Supplemental Figure 6 (yellow) using the RMSF values of 14,069 NMR models (as explained in the results section 3.5.2).

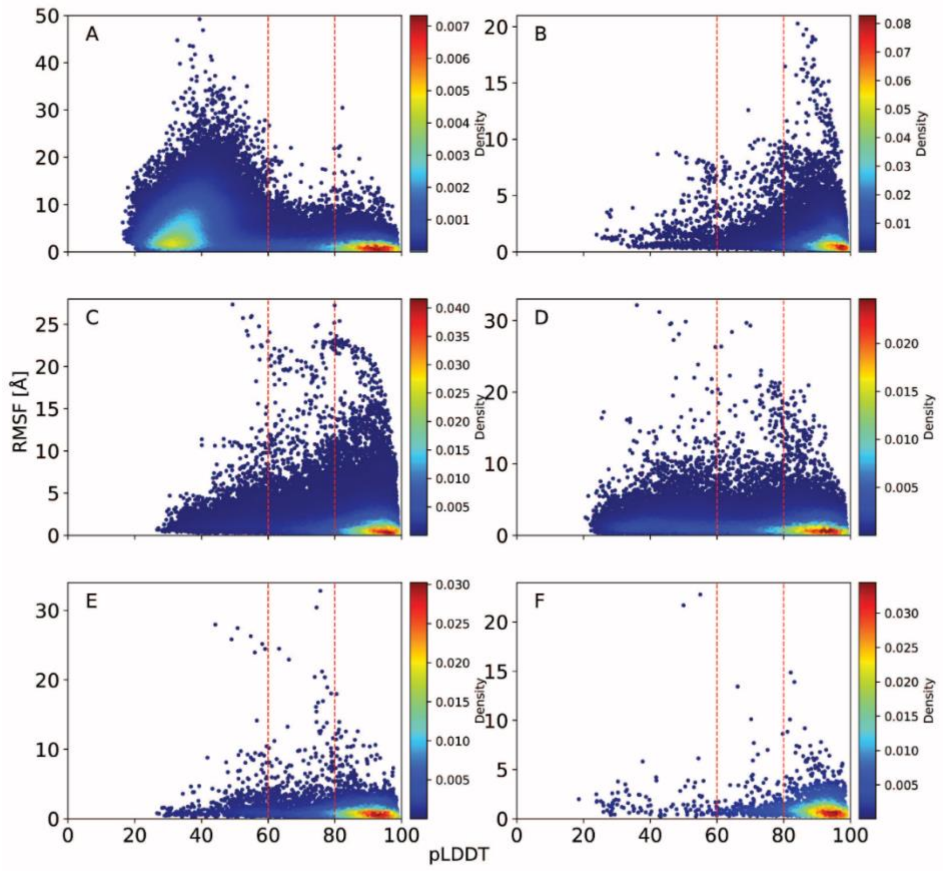


Supplemental Figure 16 **Comparison of pLDDT and RMSF.** RMSF values versus pLDDT value of each amino acid, visualized with a Gaussian kernel estimator for $S^2_{\mathrm{RCI}}$ data set. One subplot for each secondary structure element: A) coil (N = 105,172), B) strand (N = 54,786), C) $\alpha$-helix (N = 109,639), D) turn (N = 58,328), E) $3_{10}$-helix (N = 7,931), and F) bridge (N = 2,445), where N represents number of amino acid residues. The red vertical lines divide the dataset into high pLDDT ($\geq 80$), mid ($\geq 60$ pLDDT $< 80$) and low ($< 60$) pLDDT regions.

Supplemental Table 5 RMSF values are grouped according to pLDDT in low-pLDDT, mid-pLDDT and high-pLDDT. The table reports the minimum, maximum, mean, and standard deviation for each group.

| | low-pLDDT | | | | mid-pLDDT | | | | high-pLDDT | | | |
|---|---|---|---|---|---|---|---|---|---|---|---|---|
| **secondary structure** | **min** | **max** | **mean** | **std** | **min** | **max** | **mean** | **std** | **min** | **max** | **mean** | **std** |
| Coil | 0.22 | 49.24 | 5.65 | 4.43 | 0.21 | 26.71 | 1.89 | 2.18 | 0.14 | 30.48 | 1.27 | 1.29 |
| Strand | 0.24 | 8.82 | 1.89 | 1.86 | 0.21 | 12.6 | 1.35 | 1.46 | 0.14 | 20.28 | 1.11 | 1.13 |
| α-Helix | 0.25 | 27.35 | 1.87 | 1.83 | 0.18 | 27.26 | 1.84 | 2.26 | 0.13 | 25.39 | 1.38 | 1.65 |
| Turn | 0.20 | 32.16 | 2.16 | 2.03 | 0.15 | 29.68 | 1.74 | 2.16 | 0.13 | 20.95 | 1.33 | 1.36 |
| $3_{10}$-Helix | 0.25 | 27.95 | 2.08 | 3.19 | 0.21 | 32.8 | 1.89 | 2.78 | 0.16 | 17.96 | 1.14 | 1.16 |
| Bridge | 0.22 | 22.78 | 1.89 | 3.00 | 0.26 | 13.43 | 1.34 | 1.53 | 0.15 | 14.87 | 1.20 | 1.16 |

Supplemental Table 6 RMSF values are grouped according to $S^2_{RCI}$ values in flexible, ambiguous, and rigid. The table reports the maximum, maximum, mean, and standard deviation for each group.

| | flexible (< 0.70) | | | | ambiguous (0.70-0.80) | | | | rigid (>= 0.80) | | | |
|---|---|---|---|---|---|---|---|---|---|---|---|---|
| **secondary structure** | **min** | **max** | **mean** | **std** | **min** | **max** | **mean** | **std** | **min** | **max** | **mean** | **std** |
| Coil | 0.21 | 25.06 | 2.43 | 2.60 | 0.20 | 16.88 | 1.40 | 1.58 | 0.19 | 14.74 | 1.30 | 1.47 |
| Strand | 0.23 | 15.44 | 1.38 | 1.66 | 0.20 | 15.16 | 1.28 | 1.33 | 0.17 | 14.77 | 1.13 | 1.25 |
| α-Helix | 0.23 | 20.99 | 2.04 | 2.06 | 0.21 | 16.51 | 1.56 | 1.72 | 0.15 | 19.04 | 1.23 | 1.35 |
| Turn | 0.19 | 22.01 | 1.89 | 1.79 | 0.20 | 17.37 | 1.52 | 1.70 | 0.20 | 14.70 | 1.35 | 1.40 |
| $3_{10}$-Helix | 0.23 | 11.44 | 1.70 | 1.94 | 0.21 | 10.54 | 1.45 | 1.45 | 0.21 | 11.74 | 1.21 | 1.32 |
| Bridge | 0.29 | 9.20 | 1.52 | 1.49 | 0.22 | 10.09 | 1.27 | 1.34 | 0.21 | 14.87 | 1.37 | 1.73 |

Supplemental Table 7 Pearson correlation coefficients and p-values are provided for the following comparisons: RMSF and pLDDT, and RMSF and $S^2_{RCI}$. These correlations are analysed for each group within the metrics (pLDDT and $S^2_{RCI}$) as well as for the full range of metrics (including both pLDDT and $S^2_{RCI}$), with and without the classification of secondary structure elements (all SS).

| | Coil | | Strand | | α-helix | | Turn | | $3_{10}$-helix | | Bridge | | all SS | |
|---|---|---|---|---|---|---|---|---|---|---|---|---|---|---|
| | Pearson correlation coefficient | p-value | Pearson correlation coefficient | p-value | Pearson correlation coefficient | p-value | Pearson correlation coefficient | p-value | Pearson correlation coefficient | p-value | Pearson correlation coefficient | p-value | Pearson correlation coefficient | p-value |
| low-pLDDT | 0.16 | 0.00 | 0.21 | 4.62E-6 | 0.16 | 9.41E-36 | 0.07 | 9.71E-15 | 0.13 | 1.28E-3 | 0.07 | 4.99E-1 | -0.04 | 1.93E-26 |
| mid-pLDDT | -0.17 | 8.98E-55 | -0.01 | 4.51E-1 | -0.04 | 9.19E-7 | -0.04 | 1.93E-6 | -0.04 | 1.91E-1 | 0.00 | 9.63E-1 | -0.07 | 2.99E-47 |
| high-pLDDT | -0.16 | 1.90E-157 | -0.12 | 8.7E-165 | -0.10 | 1.28E-183 | -0.16 | 7.83E-208 | -0.17 | 3.16E-41 | -0.18 | 5.08E-17 | -0.13 | 0.00 |
| pLDDT | -0.43 | 0.00 | -0.12 | 1.07E-185 | -0.11 | 4.49E-319 | -0.20 | 0.00 | -0.19 | 1.08E-67 | -0.14 | 3.51E-12 | -0.24 | 0.00 |
| flexible (< 0.70) | -0.26 | 3.22E-66 | -0.11 | 4.84E-3 | -0.01 | 5.62E-1 | -0.11 | 9.5E-13 | -0.16 | 1.23E-2 | -0.14 | 1.34E-1 | -0.17 | 6.97E-75 |
| ambiguous (0.70-0.80) | -0.07 | 7.65E-5 | -0.04 | 2.57E-2 | -0.08 | 2.07E-4 | -0.03 | 5.21E-2 | -0.02 | 7.13E-1 | -0.18 | 1.44E-2 | -0.05 | 4.03E-10 |
| rigid (>= 0.80) | -0.05 | 1.27E-3 | -0.06 | 6.85E-15 | -0.05 | 2.66E-14 | -0.03 | 8.36E-3 | -0.08 | 3.34E-3 | -0.07 | 1.91E-1 | -0.06 | 3.59E-44 |
| $S^2_{RCI}$ | -0.32 | 1.06E-278 | -0.08 | 5.2E-26 | -0.15 | 4.72E-124 | -0.15 | 1.15E-76 | -0.14 | 4.67E-11 | -0.07 | 6.56E-2 | -0.22 | 0.00 |

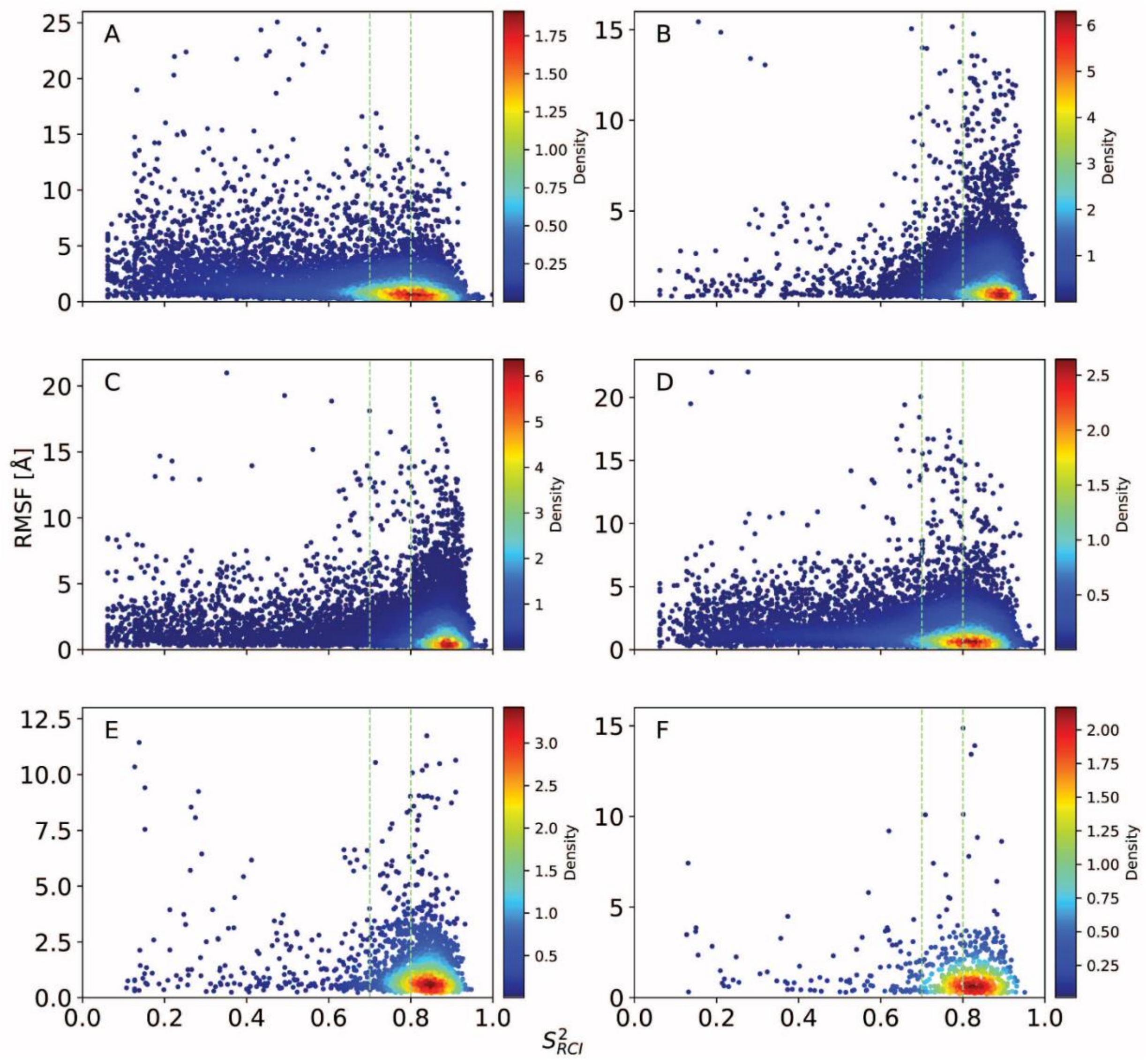


Supplemental Figure 17 **Comparison of $S^2_{\mathrm{RCI}}$ and RMSF.** RMSF values versus $S^2_{\mathrm{RCI}}$ value of each amino acid, visualized with a Gaussian kernel estimator for truncated $S^2_{\mathrm{RCI}}$ data set. One subplot for each secondary structure element: A) coil (N = 11,634), B) strand (N = 18,640), C) $\alpha$-helix (N = 25,861), D) turn (N = 14,759), E) $3_{10}$-helix (N = 2,250), and F) bridge (N = 670), where N represents number of amino acid residues. The green vertical lines divide the dataset into flexible (<0.70), ambiguous (0.70 - 0.80), and rigid ($\geq$ 80) regions.

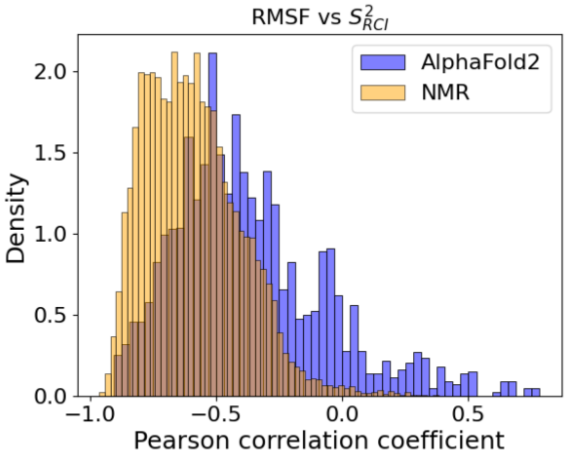


Supplemental Figure 18 **Pearson correlation coefficients between RMSF and $S^2_{\mathrm{RCI}}$** Distribution of Pearson correlation coefficients between RMSF values and $S^2_{\mathrm{RCI}}$ values of amino acids for AlphaFold2 models (blue) and NMR models (yellow).

### Examples of $S^2_{RCI}$ and RMSF correlation for Alphafold2 and NMR models

The correlation between the per-residue RMSF and per-residue $S^2_{\mathrm{RCI}}$ values of a given protein is generally expected to be *negative*, because rigid regions would correspond to high RMSF and low $S^2_{\mathrm{RCI}}$. There were a few NMR models that showed (unexpected) positive correlation between $S^2_{\mathrm{RCI}}$ and RMSF.

### Example of a protein where the AlphaFold2 model has a slightly weaker negative $S^2_{RCI}$ versus RMSF correlation than the NMR model.

The Pearson correlation coefficient between RMSF and $S^2_{\mathrm{RCI}}$ values is $-0.52$ for the AlphaFold2 model of protein Q96LL9-2YUA (BMRB id 11144). For the 20 NMR structures, 19 have a Pearson correlation coefficient between RMSF and $S^2_{\mathrm{RCI}}$ that lie in the range $-0.65$ to $-0.81$ and the remaining model shows $-0.40$. Therefore, the AlphaFold2 model has weaker correlation than most of the NMR models. In addition, in the figures below, the NMR model shows residues with ambiguous secondary structure (grey). The ambiguous residues were computed by the dumb consensus of secondary structure with a $70\%$ threshold within the NMR ensemble.

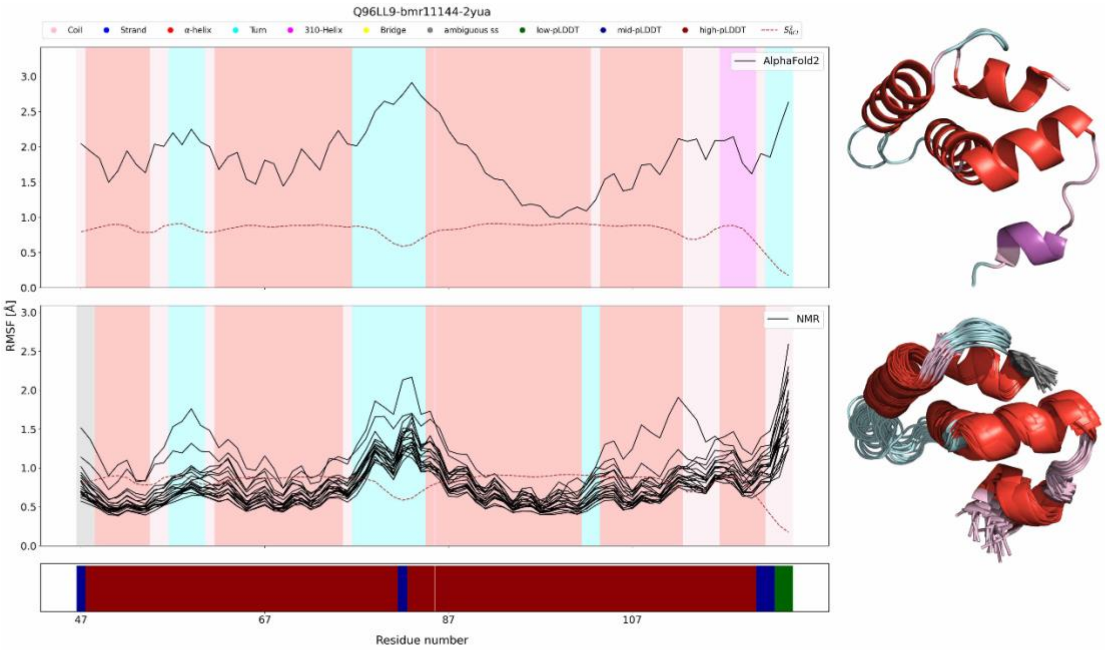


Supplemental Figure 19 **Example of a protein (Q96LL9-2YUA).** The 3D structures with secondary structure mapping colours are shown on the right: AlphaFold2 model (right top) and NMR ensemble (right bottom, 20 NMR models shown at once). Comparing $S^2_{\mathrm{RCI}}$ (red, dashed) and RMSF (black line) of the AlphaFold2 model and the NMR ensemble (for this protein, there were 20 NMR models within the ensemble). The secondary structure is indicated with shaded regions. The pLDDT of the sequence is shown below the plots. (Color legends at the top of the figure.)

#### Example of a protein where the AlphaFold2 model has a slightly stronger negative $S^2_{RCI}$ versus RMSF correlation than (most of) the NMR models.

The Pearson correlation coefficient between RMSF and $S^2_{\mathrm{RCI}}$ values is $-0.91$ for the AlphaFold2 model of protein Q922K9-2D8J (BMRB id: 11214). For the 20 NMR models within the ensemble, the Pearson correlation coefficient between RMSF and $S^2_{\mathrm{RCI}}$ lie in the range $-0.63$ to $-0.91$, where 19 models show correlation coefficient below $-0.91$. Therefore, the AlphaFold2 model has stronger negative correlation than most of the NMR models.

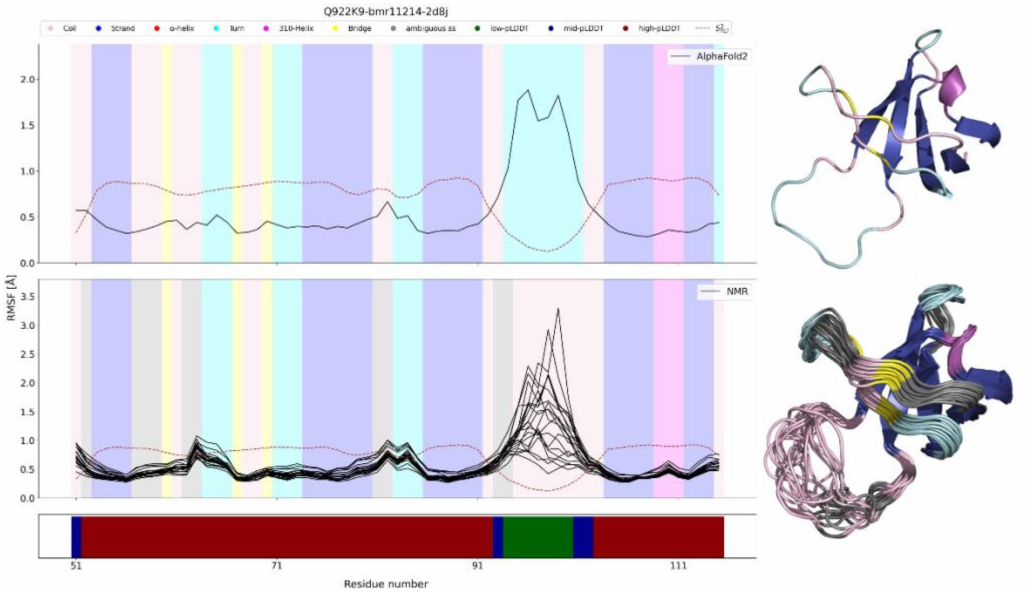


Supplemental Figure 20 **Example of a protein (Q922K9-2D8J).** The 3D structures with secondary structure mapping colours are shown on the right: truncated AlphaFold2 model (right top) and NMR ensemble (right bottom, 20 NMR models within the ensemble shown at once). Comparing $S^2_{\mathrm{RCI}}$ (red, dashed) and RMSF (black line) of the AlphaFold2 model and the NMR models (for this protein, there were 20). The secondary structure is indicated with shaded regions. The pLDDT of the sequence is shown below the plots. (Color legends at the top of the figure.)

### Example of a protein where the AlphaFold2 model has a (unexpected) positive $S^2_{RCI}$ versus RMSF correlation, while the NMR models show negative correlation.

The Pearson correlation coefficient between RMSF and $S^2_{\mathrm{RCI}}$ values is $0.63$ for the AlphaFold2 model and ranges from $-0.62$ to $-0.85$ for the $20$ NMR models within the ensemble of protein Q02053-2V31 (BMRB id 18758).

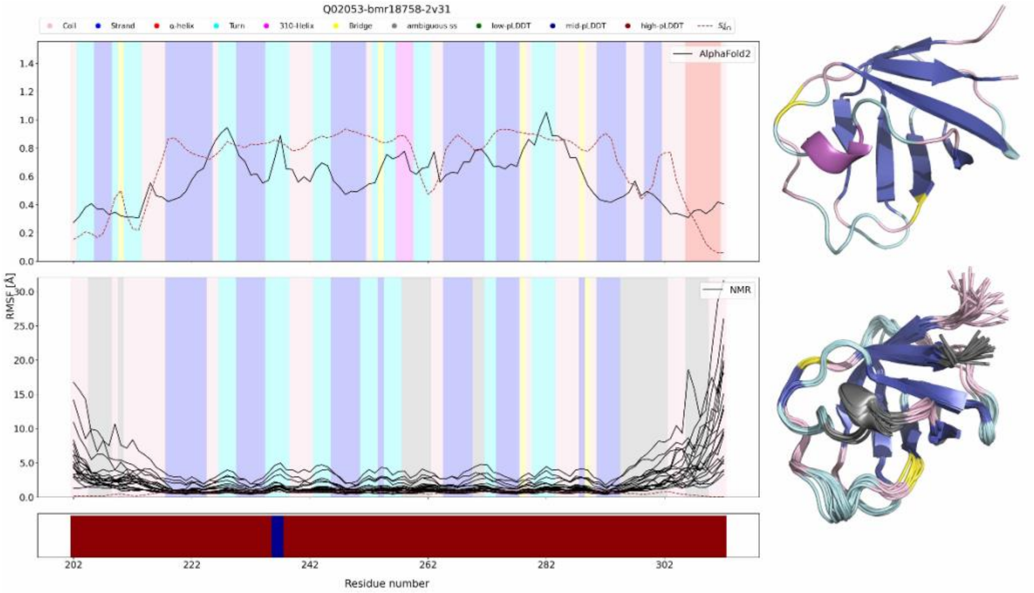


Supplemental Figure 21 **Example of a protein (Q02053-2V31).** The 3D structures with secondary structure mapping colours are shown on the right: truncated AlphaFold2 model (right top) and NMR ensemble (right bottom, 20 NMR models within the ensemble shown at once). Comparing $S^2_{\mathrm{RCI}}$ (red, dashed) and RMSF (black line) of the AlphaFold2 model and the NMR models (for this protein, there were 20). The secondary structure is indicated with shaded regions. The pLDDT of the sequence is shown below the plot. (Color legends at the top of the figure.)

#### Example of a protein where the AlphaFold2 model has a negative $S^2_{RCI}$ versus RMSF correlation, while the NMR structure shows (unexpected) positive correlation.

The Pearson correlation coefficient between RMSF and $S^2_{\mathrm{RCI}}$ values is $-0.4$ for the AlphaFold2 model and ranges from $0.00$ to $0.09$ for the 20 NMR structures of protein P37665-2N48. (BMRB id 15683).

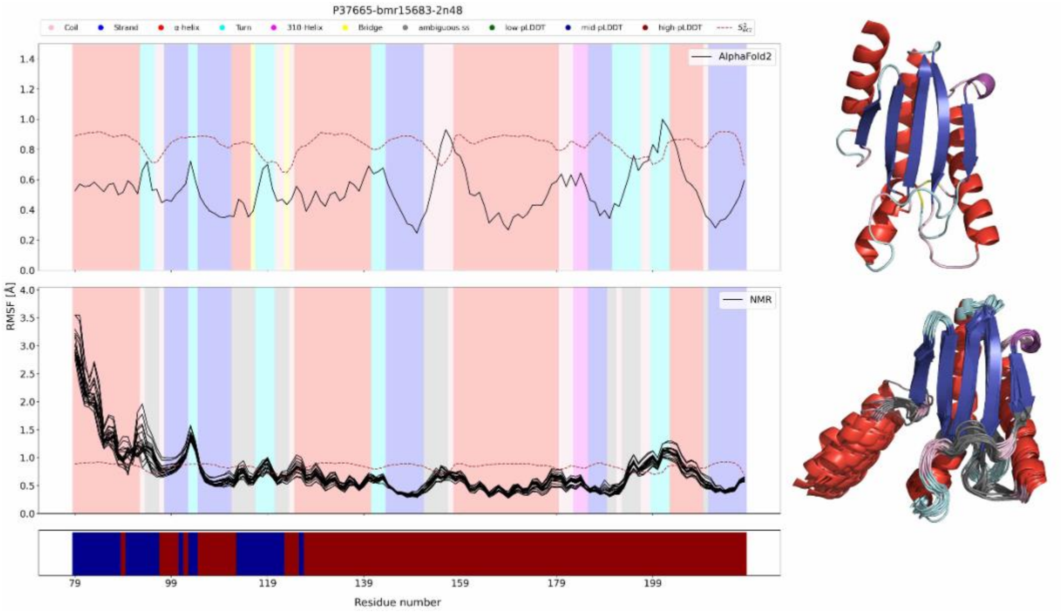


Supplemental Figure 22 **Example of a protein (P37665-2N48).** The 3D structures with secondary structure mapping colours are shown on the right: truncated AlphaFold2 model (right top) and NMR ensemble (right bottom, 20 NMR models within the ensemble shown at once). Comparing $S^2_{\mathrm{RCI}}$ (red, dashed) and RMSF (black line) of the AlphaFold2 model and the NMR models (for this protein, there were 20). The secondary structure is indicated with shaded regions. The pLDDT of the sequence is shown below the plots. (Color legends at the top of the figure.)

#### Example of a protein where the 88% of overlapping amino acid sequence between AlphaFold2 NMR shows conflicting secondary structure.

The Pearson correlation coefficient between RMSF and $S^2_{\mathrm{RCI}}$ values is $-0.55$ for the AlphaFold2 model and ranges from $-0.74$ to $-0.86$ for the 20 NMR structures of protein P0AFW0-2LCL. (BMRB id 17615).

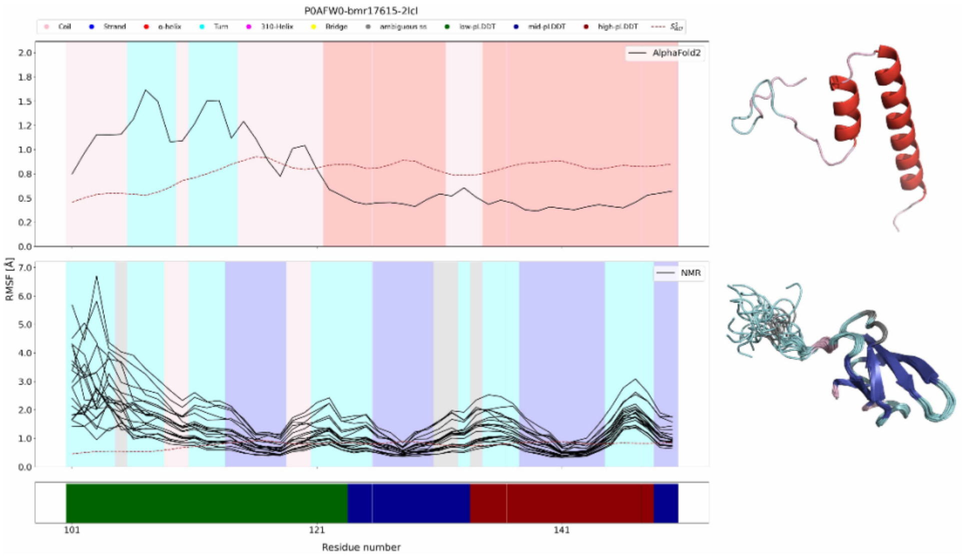


Supplemental Figure 23 **Example of a protein (P0AFW0-2LCL).** The 3D structures with secondary structure mapping colours are shown on the right: AlphaFold2 model overlapping with NMR sequence (right top) and NMR ensemble (right bottom, 20 NMR models within the ensemble shown at once). Comparing $S^2_{\mathrm{RCI}}$ (red, dashed) and RMSF (black line) of the truncated AlphaFold2 model and the NMR models (for this protein, there were 20). The secondary structure is indicated with shaded regions. The pLDDT of the sequence is shown below the plot. The sequence in the RMSF plot shows sequence from 101-150 amino acids, while the structure shows 101-161 amino acids (Color legends at the top of the figure.)

### Conflicting secondary structure elements between AlphaFold2 and NMR models

Using STRIDE, a secondary structure (SS) element is assigned to each residue of the AlphaFold2 model of a protein, and to each residue of the models in the NMR ensemble of the protein. When a residue has an equal assignment in all models (one AlphaFold2 model and one (or more) NMR models), we say that the residue has identical SS. When a residue has a different assigned SS in the AlphaFold2 model compared to its assigned SS in all the protein's NMR models, we say that the residue has a conflicting SS. Besides these residues with conflicting SS and identical SS, there is a third group of residues: a residue might have an AlphaFold2 assigned SS that is identical to the SS in some of the NMR models but conflicting in some of the other NMR models of the protein.

There are 746 unique proteins with 746 AlphaFold2 models and 746 NMR ensembles (totaling 14,069 NMR models), corresponding to 14,069 AlphaFold2-NMR pairs (see main text). Out of the 74,879 unique residues of these proteins that are present in the AlphaFold2 sequence and the NMR models (overlapping), several residues (19,561) from one or more NMR models of the same ensemble exhibit indeed both conflicting and identical secondary structures. This variability arises because different NMR models within the same ensemble can show different secondary structures for the same residues. These residues are shown as the overlap between conflicting and identical secondary structures in Supplemental Figure 24. The distribution of $S^2_{\mathrm{RCI}}$, RMSF, and pLDDT for residues with conflicting secondary structures (6,738 residues) is shown in Supplemental Figure 25. The Pearson correlation coefficient between $S^2_{\mathrm{RCI}}$ and RMSF for residues with conflicting secondary structures (SS) is -0.17 (p-value = $1.26 \times 10^{-44}$, N=6,258 where N is the number of amino acids with $S^2_{\mathrm{RCI}}$ available values). For $S^2_{\mathrm{RCI}}$ and pLDDT, the Pearson correlation is 0.44 (p-value = $0.94^{-308}$, N=6,258), and it is -0.14 (p-value = $6.96^{-35}$, N= 6,738) for RMSF and pLDDT.

We also examined the conflicting SS residues for each structure in 14,069 AlphaFold2-NMR pairs, identifying a total of 14,006 AlphaFold2-NMR pairs with conflicting SS residues. For these 14,006 pairs, we computed the difference in the Pearson correlation coefficients ($\rho^{NMR}_{k,m}$ and $\rho^{NMR}_{k,m}$, for detailed explanation see results section 3.5.2 in main) of AlphaFold2-NMR pairs Eq. 5. The values $\Delta\rho_{k,m} > 0$ indicates that $\rho^{NMR}_{k,m}$ is stronger than $\rho^{AF2}_{k,m}$ indicating, while $\Delta\rho_{k,m} < 0$ indicates that $\rho^{AF2}_{k,m}$ is stronger than $\rho^{NMR}_{k,m}$. Out of 14,006 AlphaFold2-NMR pairs, 9,994 showed stronger $\rho^{NMR}_{k,m}$, and the remaining 4,012 pairs showed stronger $\rho^{AF2}_{k,m}$. The distribution of conflicting SS residues for both cases is shown in Supplemental Figure 26. For AlphaFold2 models where correlation between $S^2_{\mathrm{RCI}}$ vs RMSF is stronger than NMR models, the percentage of conflicting SS residues range from 0.86 % to 80.48 %, with an average of 19.46

± 10.18% conflicting SS residues across the overlapping sequences of AlphaFold2 and NMR models. In comparison, for NMR models, the range is from 1.01 % to 88.00 % with an average of 19.07 ± 9.91 % conflicting SS residues.

$$\Delta\rho_{k,m} = \rho_{k,m}^{AF2} - \rho_{k,m}^{NMR} \tag{5}$$

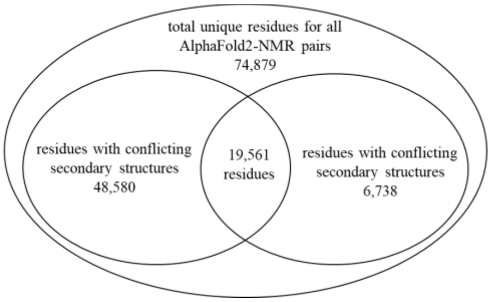


Supplemental Figure 24 **Total conflicting secondary structure residues** The Venn diagram representing total number of unique residues for 14,069 AlphaFold2-NMR pairs with conflicting secondary structure residues and identical secondary structure residues.

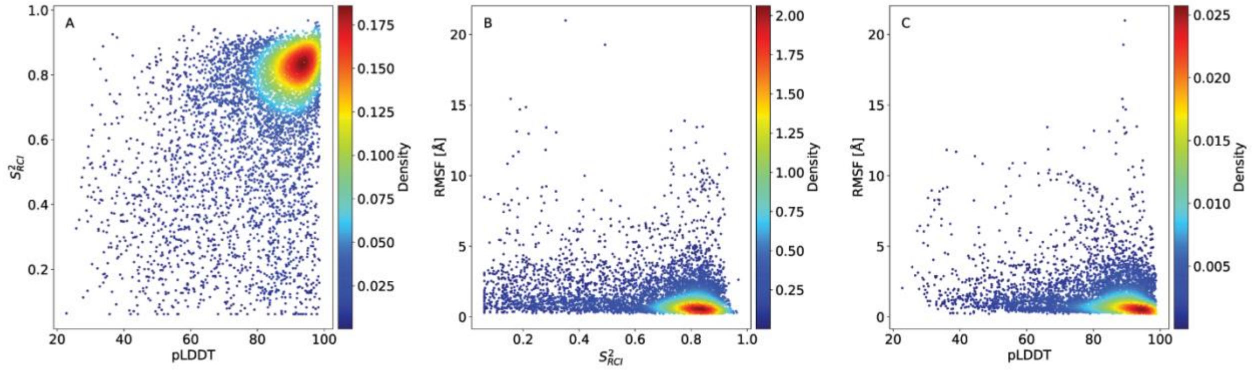


Supplemental Figure 25 **Conflicting secondary structure residues.** A) $S^2_{\mathrm{RCI}}$ vs pLDDT, B) $S^2_{\mathrm{RCI}}$ vs RMSF, and C) pLDDT vs RMSF of 6,738 residues with conflicting secondary structures between AlphaFold2-NMR pairs are shown. The A, B, and C are visualized with a Gaussian kernel estimator between their corresponding x-axis and y-axis variables.

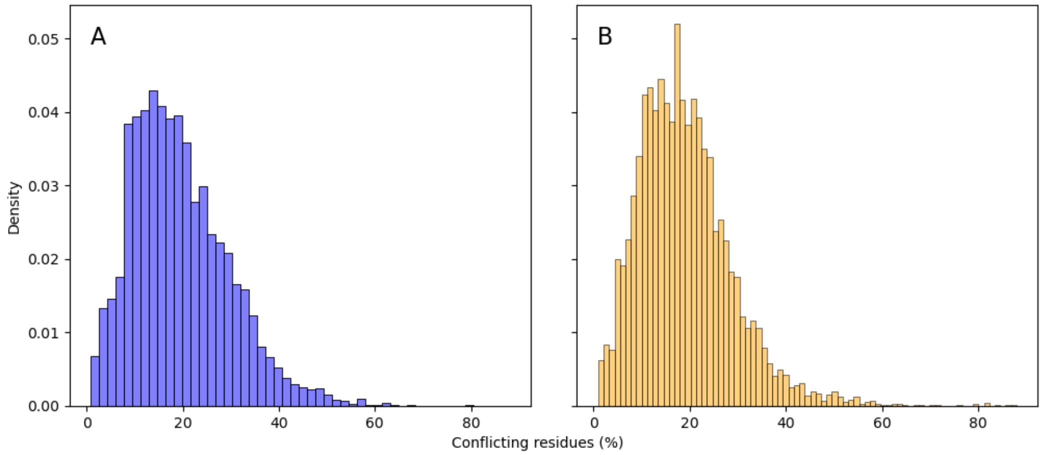


Supplemental Figure 26 **Distribution of conflicting SS residues in AlphaFold2-NMR pairs.** The distribution of conflicting SS residues is shown as percentage on x-axis for A) AlphaFold2 models where the correlation between $S^2_{\mathrm{RCI}}$ vs RMSF is stronger than NMR models, and B) NMR models where the correlation between $S^2_{\mathrm{RCI}}$ vs RMSF is stronger than AlphaFold2 models.

Supplemental Table 8 **Examples of proteins with Pearson correlation coefficients between $S^2_{\mathrm{RCI}}$ vs RMSF.** The Pearson correlation coefficients of specific proteins with their unique UniProt ID are reported for their respective AlphaFold2 and NMR models, including the number of conflicting residues occurring between the overlapping sequence of between the AlphaFold2 and NMR models, and total number of residues in the non-truncated and truncated AlphaFold2 models. For the NMR models, an example of only one structure from the NMR ensemble is provided.

| UniProt ID | correlation coefficient (AF2) | correlation coefficient (NMR) | no. of conflicting residues | total overlapping residues | conflicting residues (%) | total no. of residues (non-truncated) | total no. of residues (non-truncated) |
|---|---|---|---|---|---|---|---|
| C3VPR6 | -0.75 | -0.73 | 13.85 | 87 | 15.92 | 1915 | 1898 |
| O43157 | -0.60 | -0.46 | 21.95 | 112 | 19.60 | 2135 | 2122 |
| O60885 | -0.65 | -0.78 | 15.00 | 83 | 18.07 | 1362 | 1315 |
| P00519 | -0.89 | -0.83 | 26.50 | 97 | 27.32 | 1130 | 1081 |
| P16157 | -0.64 | -0.66 | 15.70 | 104 | 15.10 | 1881 | 1817 |
| P26039 | -0.82 | -0.78 | 12.00 | 132 | 9.09 | 2541 | 2523 |
| P35670 | -0.81 | -0.86 | 29.00 | 161 | 18.01 | 1465 | 1352 |
| P36006 | -0.84 | -0.87 | 13.05 | 69 | 18.91 | 1272 | 1230 |
| P38398 | -0.58 | -0.57 | 29.07 | 104 | 27.95 | 1863 | 1851 |
| P59046 | -0.79 | -0.68 | 28.00 | 93 | 30.11 | 1061 | 1056 |
| Q04656 | -0.73 | -0.68 | 57.15 | 181 | 31.57 | 1500 | 1436 |
| Q53SF7 | -0.52 | -0.76 | 19.15 | 79 | 24.24 | 1128 | 1041 |
| Q63HR2 | -0.54 | -0.79 | 23.55 | 114 | 20.66 | 1409 | 1375 |
| Q92625 | -0.58 | -0.76 | 26.00 | 80 | 32.50 | 1134 | 1069 |
| Q9P212 | -0.73 | -0.86 | 25.95 | 101 | 25.69 | 2302 | 1881 |
